# Early-life glucocorticoids accelerate lymphocyte count senescence in roe deer

**DOI:** 10.1101/2024.05.24.595732

**Authors:** Lucas D. Lalande, Gilles Bourgoin, Jeffrey Carbillet, Louise Cheynel, François Debias, Hubert Ferté, Jean-Michel Gaillard, Rebecca Garcia, Jean-François Lemaître, Rupert Palme, Maryline Pellerin, Carole Peroz, Benjamin Rey, Pauline Vuarin, Emmanuelle Gilot-Fromont

## Abstract

Immunosenescence corresponds to the progressive decline of immune functions with increasing age. Although it is critical to understand what modulates such a decline, the ecological and physiological drivers of immunosenescence remain poorly understood in the wild. Among them, the level of glucocorticoids (GCs) during early life are good candidates to modulate immunosenescence patterns because these hormones can have long-term consequences on individual physiology. Indeed, GCs act as regulators of energy allocation to ensure allostasis, are part of the stress response triggered by unpredictable events and have immunosuppressive effects when chronically elevated. We used longitudinal data collected over two decades in two populations of roe deer (*Capreolus capreolus*) to test whether higher baseline GC levels measured within the first year of life were associated with a more pronounced immunosenescence and parasite susceptibility. We first assessed immunosenescence trajectories in these populations facing contrasting environmental conditions. Then, we found that juvenile GC levels can modulate lymphocyte trajectory. Lymphocyte depletion was accelerated late in life when FGMs were elevated early in life. Although the exact mechanism remains to be elucidated it could involve a role of GCs on thymic characteristics. In addition, elevated GC levels in juveniles were associated with a higher abundance of lung parasites during adulthood for individuals born during bad years, suggesting short-term negative effects of GCs on juvenile immunity, having in turn long-lasting consequences on adult parasite load, depending on juvenile environmental conditions. These findings offer promising research directions in assessing the carry-over consequences of GCs on life-history traits in the wild.

## 1. Introduction

The immune system is a key component of health and viability (Schmid-Hempel 2003). It includes two interacting arms: the innate immunity that provides a fast and generalist response to pathogens and includes the inflammatory response, and the adaptive immunity that is characterised by a slower but more specific long-term protection against pathogens (Klasing 2004). The development, maintenance, and activation of the immune system is energy-demanding and can damage tissues if activation is excessive or prolonged (Klasing 2004). Consequently, when energy acquisition is limiting, the allocation to each arm of the immune system should be traded for growth, reproduction or other maintenance-related functions (Sheldon and Verhulst 1996, Lee 2006, Martin 2009, Maizels and Nussey 2013, McDade et al. 2016). Inevitably, these costs and the subsequent trade-offs can explain, at least partly, why the efficiency of the immune system cannot be maximised and maintained throughout life, and why it declines with increasing age, a process defined as ‘immunosenescence’ (Makinodan 1980). Coming mostly from human and laboratory mammal studies (Pawelec et al. 2020, but see Peters et al. 2019 for a review in free-ranging and captive animals), accumulated empirical evidence shows consistent declines of the adaptive arm and no clear changes in the innate arm, even if its inflammatory component shows consistent increase (leading to a chronic low-grade inflammatory state, named ‘*inflammageing*’, which constitutes a hallmark of the ageing process; Franceschi et al. 2000, 2007, López-Otín et al. 2023). These age-associated changes in immune functioning are generally the underlying causes of an increased susceptibility to pathogens (such as parasites) with age in mammals in the wild (Hayward et al. 2009, 2015, Froy et al. 2019, but see Hämäläinen et al. 2015), ultimately leading to higher morbidity and mortality risks and lower reproductive success (Sheldon and Verhulst 1996, Bauer and Fuente 2016, Froy et al. 2019). Accordingly, these consequences on populations make the identification of the ecological and physiological drivers of immunosenescence in the wild an important research topic.

The resource-based allocation trade-off between immunity and other biological functions (Sheldon and Verhulst 1996, McDade et al. 2016) is expected to be mediated, at least partially, by glucocorticoids (GCs). GCs are steroid hormones with pleiotropic effects. Although they are mainly known for their involvement in the stress response (Wingfield 2013, Hau et al. 2016), their primary role is to regulate the organism energy balance, and as a result, they modulate energy allocation among growth, reproduction, and somatic maintenance (Hau et al. 2016). Therefore, GCs contribute to allostasis (*i.e.* achieving stability through change) by ensuring energy homeostasis (McEwen and Wingfield 2003, 2010, Romero et al. 2009), according to predictable and unpredictable events. Unpredictable environmental perturbations elicit a stress response involving partly the activation of the hypothalamic-pituitary-adrenal (HPA) axis resulting in the release of GCs. Short-term (*i.e.* acute; within minutes to hours) GC elevation promotes reallocation of stored energy to processes enhancing survival (Sapolsky et al. 2000), after what GCs quickly return to baseline levels to support daily maintenance of energy homeostasis. Following exposure to repeated or long-lasting stressors, however, GC concentrations can remain elevated for longer periods of time (*i.e.* days to weeks). Long-term GC elevation can lead to ‘allostatic overload’, which occurs when the energy required to ensure daily and seasonal activities and to cope with the perturbations is higher than the energy intake (McEwen and Wingfield 2010). Allostatic overload can be deleterious through hypermetabolism and increased rate of telomere attrition, epigenetic age acceleration and accelerated cellular ageing (Seeman et al. 2001, Casagrande et al. 2020, Lee et al. 2021, Bobba-Alves et al. 2023), or through body growth inhibition and body condition weakening (Reeder and Kramer 2005). The detrimental consequences of allostatic overload also include the disruption of immune responses due to the immunosuppressive effects of GCs (Lee 2006, Martin 2009, Dhabhar 2014, Hau et al. 2016). Indeed, chronic GC elevations decrease the production, activity and functioning of immunoprotective cells and lead to inflammageing and immune dysregulation (Dhabhar 2014).

While chronic GC elevations have been reported to suppress immunity in laboratory studies, the carry-over consequences of GCs on immunosenescence have never been investigated in the wild. Baseline GC levels and immune parameters have effectively been experimentally shown to covary in roe deer (*Capreolus capreolus*, Carbillet et al. 2022), but longer term relationships between GCs and immunity have not yet been assessed. Similarly, GCs and parasite burden are generally positively linked on the short term in mammals (Defolie et al. 2020). The underlying mechanisms of these associations could involve the impairing effects of GCs on host immunity and their supporting role in parasite development (Herbert et al. 2022), but are not fully elucidated. While the immunosuppressive effects of GCs in chronically stressed individuals (Lee 2006, Martin 2009, Dhabhar 2014) and the consequences of acute stress-induced GC concentrations on individual performance (leading to the concept of ‘*emergency life-history stage*’, Wingfield et al. 1998) have been investigated quite intensively using laboratory animal models, the consequences of elevated baseline GC levels (resulting from repeated or chronic stress exposure) on life-history traits are more equivocal (*e.g.* Pride 2005, Pauli and Buskirk 2007, Cabezas et al. 2007, Wey et al. 2015). Similarly, the literature on stress exposure or chronically elevated GC levels and immunosenescence suggests the predominance of negative relationships (Bauer 2005, 2008, Garrido et al. 2022), but most studies so far focused on humans or laboratory rodents, and none has been performed in the wild. Importantly, these studies generally investigate covariation patterns between chronic stress exposure or chronically elevated GC levels and changes in immune parameters in elderly, rather than to the study of early-life stress and carry-over consequences in immune functioning.

Here, we took advantage of an exceptionally detailed long-term individual-based monitoring based on a capture-mark-recapture (CMR) program in two roe deer populations in the wild. This article follows-up on previous studies describing immunosenescence patterns in these populations and had two objectives: *i*) to assess and update mean individual immunosenescence patterns for 12 immune traits encompassing both adaptive and innate immune components, and for susceptibility to 4 parasites frequently occurring in roe deer based on an extended dataset and *ii*) to explore the long-term effects of juvenile GC levels on immunosenescence patterns. Previous studies in these populations showed that immunosenescence patterns were mainly driven by population-level characteristics (Cheynel et al. 2017). However, whether and how physiological markers, and more particularly GCs, modulate those patterns have not been investigated. Thus, we first assessed immunosenescence patterns and parasite trajectories accounting for confounding factors that are known to modulate mammalian senescence patterns such as current environmental context (*e.g.* Tidière et al. 2016, Cheynel et al. 2017), sex (*e.g.* Brooks and Garratt 2017, Lemaître et al. 2020) or natal environmental conditions (*e.g.* Nussey et al. 2007, Cooper and Kruuk 2018). Then, we used baseline GC levels (estimated through faecal glucocorticoid metabolites, FGMs) measured during the first year of life to investigate whether and how juvenile GCs influenced immunosenescence patterns. The juvenile stage is a major determinant of future individual performance in this income breeder (Gaillard et al. 2003) and early-life stress is expected to have detrimental consequences on adult performance (Monaghan and Haussmann 2015). We thus expected individuals exhibiting higher levels of GCs during the juvenile stage to display earlier and/or accelerated immunosenescence due to the role of GCs in resource allocation. High GCs in juveniles were also expected to result in impaired immunity and enhanced parasite development during late life (Herbert et al. 2022).

## 2. Material and methods

### 2.1. Study populations

Two forest roe deer populations at Trois-Fontaines (TF - 1360 ha) in north-eastern France (48°43’N, 4°55’E) and at Chizé (CH - 2614 ha) in western France (46°05’N, 0°25’W) have been monitored through an intensive CMR program every winter since 1975 and 1977, respectively. At TF, forest productivity is high due to rich soils and the habitat is homogeneous at a large spatial scale and of good quality. By contrast, at CH, poor soils and frequent summer droughts result in overall low forest productivity and heterogeneous habitat quality with three main tree stands (Pettorelli et al. 2006), which makes this forest less suitable for roe deer. In both populations, quality of natal environmental conditions can be measured through the cohort-specific average mass of weaned offspring at the onset of winter (*i.e.* at about 8 months of age) corrected for the Julian date of capture (Gaillard et al. 1996). This metric of environmental quality allows accounting for the influence of environmental conditions early in life on roe deer performance, which predominate over current environmental conditions in roe deer (Pettorelli et al. 2002), and captures more reliably the effect of ‘cohort quality’ (*sensu* Gaillard et al. 2003 and see Lalande et al. 2023 and Cambreling et al. 2023 for case studies for GCs-body mass relationships and senescence of antler size, respectively). Such an integrative metric of natal environmental conditions encompasses variations in early environmental conditions (*e.g.* climate, resource availability and quality, roe deer density) as well as information about individual quality (*e.g.* mother condition and/or maternal care, themselves partly dependent on environmental quality; Toïgo et al. 2006). Cohort quality was included as a categorical variable with two levels (poor *v.* good cohorts) split around the median cohort quality in each population. Captures take place each year during 10-12 days, spread from December (at TF) or January (at CH) to early March (Gaillard et al. 1993). Yearly population size estimated from CMR analyses indicate that both populations were much lower during the present study period (2010-2022, Table S1) than the maximum density recorded throughout the forty year-long monitoring. Since 2010, faecal matter is collected rectally and blood sampled from the jugular vein (1 mL/kg to a maximum of 20 mL) on weighed (to the nearest 0.1 kg and 0.05 kg in TF and CH, respectively) individuals of known sex and age (*i.e.* captured within their first year of life, either as newborn in spring, or as 8 months of age in winter).

### 2.2. Faecal Glucocorticoid Metabolites

In the present study, baseline HPA activity was estimated through faecal glucocorticoid metabolites (FGMs, Palme 2019) on juvenile roe deer (*i.e.* 8-9 months old). FGMs represent an integrative measure of GC concentrations and thus adrenocortical activity several hours before sampling (*i.e.* baseline stress; Palme, 2019), with a delay of 12 hours on average in roe deer (ranging from 6 to 23 hours, Dehnhard et al. 2001). Thus FGMs are widely used as a proxy for evaluating the baseline activity of the HPA axis of an individual (Palme 2019), notably in roe deer (Dehnhard et al. 2001, Möstl et al. 2002, Zbyryt et al. 2018).

Extraction of FGMs consisted in 0.5 g (± 0.005) of fresh faeces vortexed in 5 mL of 80% methanol and centrifuged for 15 minutes at 2500 *g* (Palme et al. 2013). The amount of FGMs was determined in an aliquot of the supernatant after a 1+9 dilution with assay buffer and was performed using a group-specific 11-oxoaetiocholanolone enzyme immunoassay (EIA), according to a method previously described in Möstl et al. (2002) and validated for roe deer (Zbyryt et al. 2018). Measurements were done in duplicate and intra- and inter-assay coefficients of variation were lower than 10% and 15%, respectively. Results of FGMs are expressed as nanograms per gram of wet faeces (ng/g).

Faeces collection in the wild is subject to several issues that might impact FGM levels. In both populations, faeces were collected rectally and immediately frozen at −80 °C after collection, except prior to 2017 in TF where faeces were stored at 4 °C and frozen at −20 °C within 24 hours. The time from faecal collection to freezing can impact FGM levels due to different bacterial activity (Lexen et al. 2008, Hadinger et al. 2015, Carbillet et al. 2023b). Secondly, FGMs are likely to show variation throughout the capture period due to seasonal variation in HPA axis activity (Bubenik et al. 1998, Huber et al. 2003). Finally, animals are usually manipulated and sampled from 1 to 4 hours following the capture, but this delay can overpass 6 hours in exceptional cases so that the variation in the time delay between capture and sampling can also impact FGM levels. We first tested whether these three protocol issues affected the FGM levels measured in 162 individuals during their first winter. A linear mixed effect model (LMM) accounting for *i)* the Julian date of capture (linear and quadratic), *ii)* the time delay (min) between capture and sampling, and *iii)* whether faeces were immediately frozen at −80 °C or frozen at −20 °C within 24 hours, showed no effect of these confounds on log-transformed values of FGMs (Table S2). During the capture period, roe deer’s diet is mostly composed of brambles (*Rubus* sp.) and ivy (*Hedera helix*) and similar in both study sites (Tixier and Duncan 1996), so we do not expect any bias in the measurement of FGM between populations that could be associated with diet.

### 2.3. Immune traits

To assess immunosenescence in both the innate and adaptive components of the immune function, we monitored 12 immune traits of both the humoral and cellular immune activity (Cheynel et al. 2017) measured between 2010 and 2021 (as well as 2022 for haemagglutination (HA) and haemolysis (HL) scores). In the field, blood samples are partitioned upon collection into *i*) EDTA tubes for white blood cells (WBCs) count and *ii*) serum collection tubes which were centrifuged 10 minutes at 3000 *g* before the serum is collected and frozen for subsequent HA-HL and globulin measurements.

#### 2.3.1. Cellular innate immunity

Cellular innate immunity was assessed by describing the composition of the WBC population. The first hundred WBCs are counted in Wright-Giemsa-stained blood smears to estimate the proportions of each cell type. Then proportions are combined with the total WBC count to obtain concentrations of the different types of WBCs (10^3^ cells/mL) (Houwen 2001). WBC population is composed of five different cell types: neutrophils are the most numerous WBCs and, together with monocytes, are key components of the innate arm of the immune system through their antimicrobial/biotoxic and phagocytic activity. Moreover, neutrophils and monocytes have recently been identified as having a role in auto-immune and inflammatory diseases when their activation is dysregulated (Auffray et al. 2009, Burn et al. 2021). Basophils, the least abundant WBCs, have a specific role in the inflammatory response as they are particularly active at sites of ectoparasite infections such as ticks, and communicate with the adaptive arm of the immune system (Siracusa et al. 2010, Karasuyama et al. 2011). Eosinophils are also involved in the inflammatory responses and reaction to allergies alongside basophils and mast cells. They also play a role in the defence against internal parasites such as helminths, and in the modulation of both components of immunity. If dysregulated, their cytotoxic action can have detrimental consequences on the organism (Rothenberg and Hogan 2006). Finally, lymphocytes are involved in the cellular adaptive immunity, as described below.

#### 2.3.2. Humoral innate immunity

To evaluate the humoral part of the innate immune system, we performed HA-HL assays, as described in Matson et al. (2005) and previously used on roe deer blood samples (Gilot-Fromont et al. 2012, Cheynel et al. 2017, Carbillet et al. 2022, 2023a). Briefly, haemagglutination [HA, 10^−2^ log(dilution)] measures the plasmatic ability of an individual to agglutinate exogeneous cells, as a proxy of the concentration of circulating natural antibodies (NAbs). Similarly, haemolysis [HL, 10^−2^ log(dilution)] measures the plasmatic activity of the complement, a group of proteins acting in chain reactions to provoke the lysis of exogeneous cells in the presence of antigen-antibody complexes (Matson et al. 2005). Secondly, we assessed the concentrations (mg/mL) of alpha1-globulins, alpha2-globulins and beta-globulins to characterise the level of inflammatory proteins involved in the acute phase response, a part of the early innate immune system (Cray et al. 2009). It was done using refractometry and automatic agarose gel electrophoresis (HYDRASYS, Sebia, Evry, France) that separates blood proteins into five fractions: albumin, alpha1-, alpha2-, beta- and gamma-globulin fractions. More specifically, we also assayed haptoglobin concentration, a protein from the alpha2-globulin fraction produced in case of chronic inflammation and infection. Haptoglobin concentration (mg/mL) was measured with a Konelab 30i automaton (Fisher Thermo Scientific, Cergy-Pontoise, France) using phase haptoglobin assay chromogenic kit (Tridelta Development LTD, County Kildare, Ireland).

#### 2.3.3. Cellular adaptive immunity

Cellular adaptive immunity was characterised by measuring the concentration of lymphocytes (10^3^ cells/mL). Lymphocytes include both B and T cells. T lymphocytes have many functions, among which the recognition of exogeneous or infected cells and can provoke cell death. B lymphocytes are an essential part of the humoral adaptive immunity, being the main antibody-producing cells. Lymphocytes also include natural killer (NK) cells, as part of the innate immunity but are a minority among lymphocytes (*e.g.* 0.5-10 % in cattle; Kulberg et al. 2004).

#### 2.3.4. Humoral adaptive immunity

Humoral adaptive immunity was measured through the concentration (mg/mL) of gamma-globulins (or immunoglobulins) which represent the majority of circulating antibodies. Gamma-globulins levels were also determined from electrophoresis as described above.

### 2.4. Parasite abundance

We counted the number of faecal propagules of 4 parasites commonly occurring in roe deer, between 2010 and 2021, as it is a reliable estimator of the parasitic load in roe deer in these populations during the capture period (Body et al. 2011). Investigated parasites encompassed pulmonary nematodes (protostrongylids), gastro-intestinal (GI) nematodes (GI strongyles and *Trichuris* sp.) and GI protozoan (coccidia, *Eimeria* sp.). For GI nematodes and protozoan, we obtained the counts of eggs per gram (EPG) and oocysts per gram (OPG) of faeces, respectively, using a modified McMaster protocol (Raynaud 1970) with a solution of magnesium sulfate (MgSO_4_, s.g = 1.26, 2010-2013), zinc sulfate (ZnSO_4_, s.g. = 1.36, 2014-2020) and saturated salt (NaCl, s.g = 1.20, 2021) and by counting in the whole chambers of the McMaster slide (*i.e.* quantitative examination with a theoretical sensitivity of 15 EPG/OPG). We also centrifuged a 14 mL tube with the remaining solution, covered with a coverslip, before seeking for the presence of parasite propagules not detected on the McMaster slide (*i.e.* control slide, qualitative examination). For pulmonary nematodes, we obtained the count of stage 1 larvae per gram of faeces (LPG) using the Baermann faecal technique (Baermann 1917).

### 2.5. Statistical analyses

All analyses were performed using R version 4.3.0 (R Core Team 2023).

#### 2.5.1. Immunosenescence and age-specific changes in parasite abundance

For the 12 immune traits and the 4 parasites, we tested different ageing trajectories using linear mixed models (LMMs): no age effect, a linear effect of age, a quadratic effect, and threshold trajectories. We tested three threshold models differing by whether pre- and post-threshold trajectories were set as constant or not: the first model corresponded to no age-related variation until a threshold age and a linear change of traits beyond that threshold, the second corresponded to a linear change of traits before the threshold age and no age-related variation beyond, and the third corresponded to two different linear changes of traits before and beyond the threshold age. Threshold models were implemented following an approach previously described and commonly used in ageing research (*e.g.* Douhard et al. 2017, Briga et al. 2019). Briefly, thresholds varied between 3 and 10 years old with one-year increments and we identified the best-fitting threshold as described in subsection 2.5.2 below. Thresholds were not tested above 10 years old because it is the last age for which we have observations of at least one individual for each population-sex-cohort group.

Immunosenescence and parasite abundance trajectory models included each immune or parasitic trait as the response variable. Parasite abundances were log-transformed as *ln(n+1)* to reach a normal distribution. For HL scores, we used the residuals of the regression of the score on the serum coloration as a response variable, to correct for the ability of the complement to cause haemolysis according to the level of haemolysis of the serum.

We expected immunosenescence patterns to differ between populations and between sexes (Cheynel et al. 2017). Early-life environmental conditions may also influence sex-specific immunosenescence because cohort-specific environmental quality has been shown to drive differential variation in adult phenotypes between males and females in the two populations (Douhard et al. 2013, Garratt et al. 2015). Models included the additive fixed effects of age (linear, quadratic or threshold function), population, sex and cohort quality, and the two-way interactions between age terms and the three other factors. Models also included four possible confounding variables as fixed effects: 1) the body mass (hereafter named ‘mass’: standardised for the Julian date of capture for juveniles; Douhard et al. 2017) as body condition affects immune phenotype (Gilot-Fromont et al. 2012) and as parasitism is negatively linked to body mass in roe deer (Body et al. 2011); 2) the age at last observation to account for the selective disappearance of individuals with low immune performance, which could ultimately lead to an under-estimation of the intensity of senescence (hereafter named ‘age-last’; van de Pol and Verhulst 2006, van de Pol and Wright 2009); 3) the Julian date of capture to account for potential temporal variations in immune parameters and parasite exposure (hereafter named ‘julian’); and 4) the time delay between capture and blood sampling, for immune but not parasitic traits, to account for changes in immune parameters in response to hormonal changes triggered by the stress of capture (hereafter named ‘delay’). We refer to these four additional fixed effects as ‘confounding effects’ hereafter. Finally, all models included random effects of the individual identity (‘ID’) to account for repeated measurements on the same individuals, as well as the year of capture nested within the population, to account for year-to-year variation in demographic and environmental conditions in both populations. At the end, we thus compared six models (*i.e.* no age effect, linear, quadratic, and three threshold age functions) per immune or parasitic trait (see Table S3 for the full list of models and their selection). We did not include random slopes of age terms as they resulted in perfect correlations between the random intercepts and slopes (Harrison et al. 2018). The normality of the residuals and the presence of outliers were visually assessed through histograms.

#### 2.5.2. Model selection for senescence trajectories

To select for the most relevant ageing trajectories, we performed model selection to compare the constant, linear, quadratic and threshold models based on the second-order Akaike Information Criterion corrected for small sample size (AICc) as implemented in the package ‘MuMIn’ (Bartoń 2021). Briefly, among the set of models within 2 ΔAICc, we considered the simplest model (*i.e.* with the lowest number of parameters) as the supported model to satisfy parsimony rules (Burnham and Anderson 2002, see also Arnold 2010; Table S3). Note that for all threshold models, for comparison with other model types, we added one parameter and 2 to their AICc values to penalise these models for the extra-parameter of the threshold, but which is not explicitly included in the models.

#### 2.5.3. Juvenile-stage FGMs and immunosenescence

We then tested whether FGMs levels measured in the juvenile period modulated the retained immunosenescence patterns by testing the effect of juvenile-stage FGMs on the rates of senescence of each immune trait. Because having individuals with both FGMs measured during their first year of life and adult immune/parasite traits substantially decreased sample sizes, we kept the structure of the previously found immunosenescence patterns and added the additive effect of FGMs. We also added the two-way interactions between FGMs and *i*) the different retained age terms and *ii*) population, sex and/or cohort quality when they were initially retained. If no detectable age change occurred in the previously retained trajectories, we only added the additive effect of early-life FGMs on the immune/parasitic traits, in addition to the confounding variables (*i.e.* ‘mass’, ‘age-last’, ‘julian’, ‘delay’) that were previously retained. Model selection tables for each trait are presented in Supplementary Information (SI, Table S4). Note that for some variables, confidence intervals of estimates might overlap 0. This is because these variables have previously been retained when describing immunosenescence patterns on the full dataset, and this structure was considered as fixed when testing FGM effects on immunosenescence rates with the reduced dataset. The random effect structure was also kept as described above, except for alpha1- and alpha2-globulin for which the inclusion of ID resulted in singularities. Therefore, when testing for the effect of FGMs during the juvenile stage on the immunosenescence rates of these two traits, we included only one observation per individual, keeping the observation for which the age at measurement was the closest to the mean age of all individuals that were sampled only once.

Once the trajectory had been identified for the 12 immune and the 4 parasitic traits, we used the same model selection approach to retain or not the additive and interacting effects of juvenile FGMs with the age function terms, and population, sex, and cohort quality factors (Table S4).

## 3. Results

### 3.1. Immune traits

Immunosenescence patterns were determined on 1021-1201 observations obtained from 483-556 individuals, depending on the investigated trait (see Table 1). Immunosenescence patterns are displayed in Figure 1, whereas immune traits for which no age variation was detected are displayed in Supplementary Information (Figure S1).

**Figure 1.**
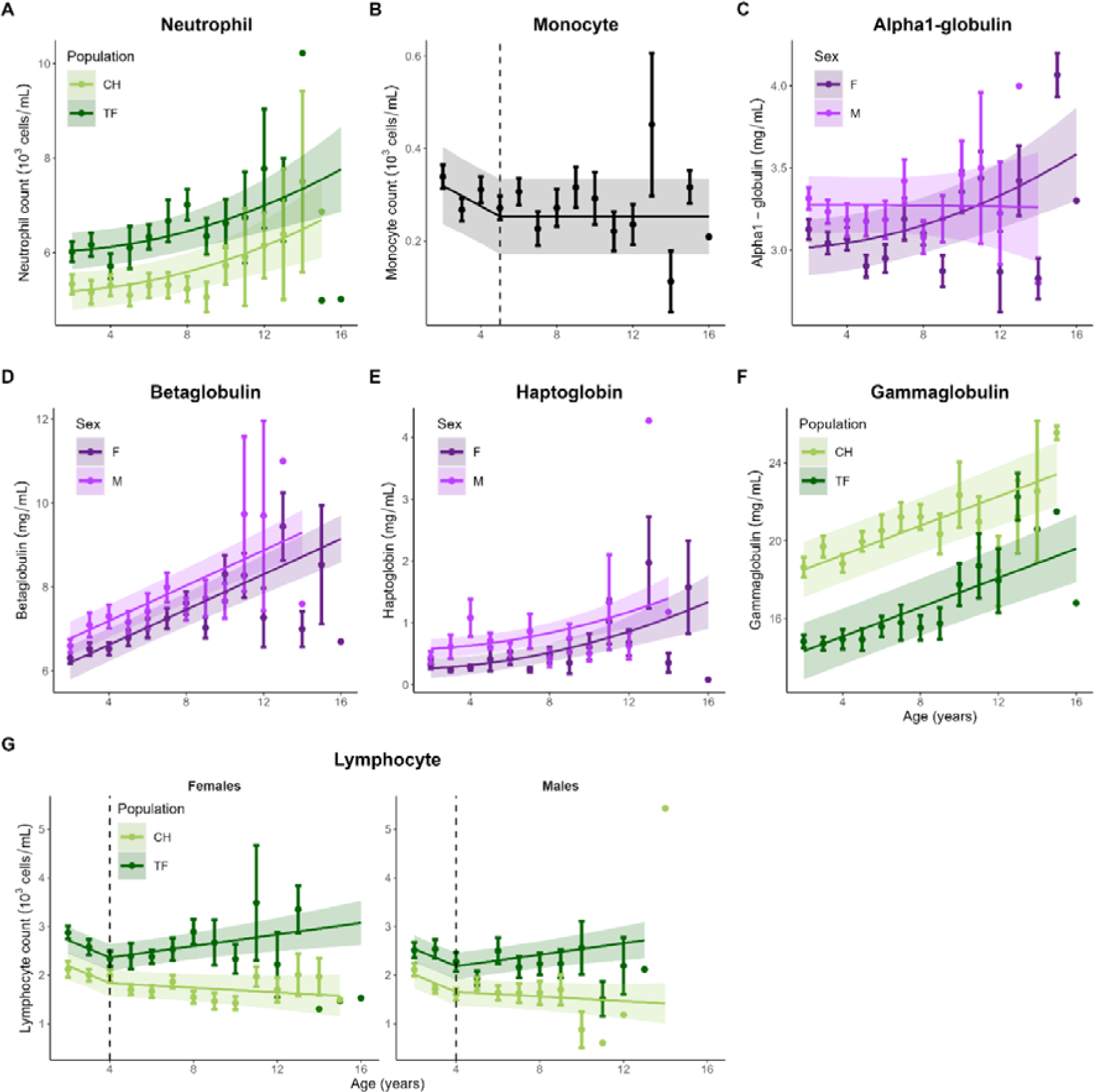
Immunosenescence patterns in roe deer. Lines are retained model predictions for each immune trait (excluding retained confounding variables for graphical representation) and shaded areas are 95% CIs. Points are age-specific average trait values ± standard errors. Dots without error bars correspond to unique observations.

**Table 1.** Selected linear mixed effect models for the 12 immune traits according to age (linear, quadratic, threshold), population (‘Pop’, reference level is Trois-Fontaines; CH: Chizé) and sex (‘Sex’, reference level is female; M: male). Models were selected based on the Akaike Information Criteria corrected for small sample sizes (AICc) as described in section 2.5.2. AICc values of retained models can be found in Tables S3. Models accounted for the following potential confounding effects: body mass (‘Mass’), age at last observation (‘Age-last’), time delay between capture and sampling (‘Delay’) and Julian day of capture (‘Julian’). All models included the random effects of the individual identity and the year of capture nested within the population. SE: standard error, R²m – R²c: marginal – conditional model variance, respectively.

| Immune trait | Model | Fixed effects | Estimate | SE | R<sup>2</sup>m | R<sup>2</sup>c |
| --- | --- | --- | --- | --- | --- | --- |
| INNATE IMMUNE SYSTEM |  |  |  |  |  |  |
| Neutrophil<br>(N = 1021 obs., n = 484 ind.) | Quadratic | Intercept | 4.81 | 0.26 | 0.09 | 0.52 |
|  |  | Age<sup>2</sup> | 0.07.10<sup>-1</sup> | 0.02.10<sup>-1</sup> |  |  |
|  |  | Pop(CH) | -0.79 | 0.26 |  |  |
|  |  | Delay | 0.04.10<sup>-1</sup> | 0.06.10<sup>-2</sup> |  |  |
| Monocyte<br>(N = 1021 obs., n = 484 ind.) | Threshold<br>(5 years of age) | Intercept | 0.25 | 0.04 | 0.01 | 0.41 |
|  |  | Age.1 | -0.02 | 0.07.10<sup>-2</sup> |  |  |
| Basophil<br>(N = 1021 obs., n = 484 ind.) | - | Intercept | 0.08 | 0.01 | 0.01 | 0.30 |
|  |  | Delay | 0.08.10<sup>-3</sup> | 0.03.10<sup>-3</sup> |  |  |
| Eosinophil<br>(N = 1021 obs., n = 484 ind.) | Sex | Intercept | 0.26 | 0.02 | 0.09 | 0.19 |
|  |  | Sex(M) | -0.04 | 0.01 |  |  |
|  |  | Delay | -0.04.10<sup>-2</sup> | 0.05.10<sup>-3</sup> |  |  |
| Haemagglutination<br>(N = 1201 obs., n = 556 ind.) | - | Intercept | 3.90 | 0.15 | - | 0.38 |
| Haemolysis<br>(N = 1191 obs., n = 555 ind.) | - | Intercept | -0.16 | 0.18 | 0.04.10<sup>-1</sup> | 0.49 |
|  |  | Delay | 0.08.10<sup>-2</sup> | 0.03.10<sup>-2</sup> |  |  |
| INFLAMMATORY SYSTEM |  |  |  |  |  |  |
| Alpha1-globulin<br>(N = 1029 obs., n = 485 ind.) | Quadratic | Intercept | 4.07 | 0.19 | 0.06 | 0.46 |
|  |  | Age<sup>2</sup> | 0.04.10<sup>-1</sup> | 0.07.10<sup>-2</sup> |  |  |
|  |  | Sex(M) | 0.31 | 0.05 |  |  |
|  |  | Age<sup>2</sup>:sex(M) | -0.02.10<sup>-1</sup> | 0.01.10<sup>-1</sup> |  |  |
|  |  | Mass | -0.04 | 0.01 |  |  |
|  |  | Julian | -0.03.10<sup>-1</sup> | 0.01.10<sup>-1</sup> |  |  |
|  |  | Age-last | -0.02 | 0.01 |  |  |
| Alpha2-globulin<br>(N = 1029 obs., n = 485 ind.) | Sex | Intercept | 5.83 | 0.15 | 0.01 | 0.23 |
|  |  | Sex(M) | -0.28 | 0.10 |  |  |
| Beta-globulin<br>(N = 1029 obs., n = 485 ind.) | Linear | Intercept | 7.70 | 0.55 | 0.12 | 0.41 |
|  |  | Age | 0.23 | 0.02 |  |  |
|  |  | Sex(M) | 0.69 | 0.13 |  |  |
|  |  | Mass | -0.10 | 0.03 |  |  |
| Haptoglobin<br>(N = 1034 obs., n = 483 ind.) | Quadratic | Intercept | 0.25 | 0.08 | 0.03 | 0.20 |
|  |  | Age<sup>2</sup> | 0.04.10<sup>-1</sup> | 0.01.10<sup>-1</sup> |  |  |
|  |  | Sex(M) | 0.32 | 0.08 |  |  |
| ADAPTIVE IMMUNE SYSTEM |  |  |  |  |  |  |
| Gamma-globulin<br>(N = 1029 obs., n = 485 ind.) | Linear | Intercept | 17.66 | 1.49 | 0.23 | 0.63 |
|  |  | Age | 0.42 | 0.05 |  |  |
|  |  | Pop(CH) | 3.70 | 1.04 |  |  |
|  |  | Mass | -0.19 | 0.06 |  |  |
| Lymphocyte<br>(N = 1021 obs., n = 484 ind.) | Threshold<br>(4 years of age) | Intercept | 3.42 | 0.20 | 0.14 | 0.48 |
|  |  | Age.1 | -0.19 | 0.04 |  |  |
|  |  | Age.2 | 0.06 | 0.02 |  |  |
|  |  | Pop(CH) | -0.55 | 0.19 |  |  |
|  |  | Sex(M) | -0.18 | 0.07 |  |  |
|  |  | Age.2:pop(C H) | -0.08 | 0.03 |  |  |
|  |  | Delay | -0.01.10<sup>-1</sup> | 0.03.10<sup>-2</sup> |  |  |

For innate traits, neutrophil levels differed between populations. Neutrophils were higher at TF, the good-quality habitat, than at CH, the low-quality habitat. The retained model evidenced a similar increase with age in both populations, without any detectable between-sex or among-cohort differences (Table 1, Figure 1A). Monocytes decreased from 2 to 5 years of age. Afterwards our approach could not detect any age-related change until the end of life (Table 1, Figure 1B). Basophils, eosinophils, and HA and HL scores did not display any detectable age-related variation, although eosinophil levels were higher in females than in males at all ages, without any evidence of between-population or among-cohort differences (Table 1, Figure S1A-D).

For the inflammatory response, alpha1-globulins, beta-globulins and haptoglobins showed an overall increase in their concentrations throughout lifespan (Table 1, Figure1C-E, Figure S1E), except in males for alpha1-globulins that were overall independent of age (Table 1, Figure 1C). Note that for alpha1-globulins, concentration in females was lower than those of males at the beginning of life but reached that of males at around 10 years of age before keeping increasing until the end of life. Beta-globulin concentration increased linearly with age and similarly in both sexes, with higher levels in males than in females, and without any between-population or among-cohort differences (Table 1, Figure 1D). Haptoglobin concentration increased quadratically across life similarly in both sexes but with higher concentration for males compared to females (Table 1, Figure 1E). We detected no age-related variation in alpha2-globulin concentration throughout life but the concentration was slightly higher in females compared to males (Table 1, Figure S1E).

Models for adaptive immunity showed that gamma-globulin concentration increased linearly throughout life and similarly in both populations, with a higher concentration in CH than TF (Table 1, Figure 1F). Lymphocyte concentration decreased in the first part of life (until 4 years of age) in both populations. Afterwards, lymphocyte count kept slightly decreasing in CH until the end of life, whereas they increased in the good-quality habitat of TF. Lymphocyte count was consistently higher at TF than at CH, and higher in males than in females (Table 1, Figure 1G). Note that the retained variables explained more variation in the adaptive immune parameters than they did for innate or inflammatory traits (see R² in Table 1). Immunosenescence onsets and rates for each trait are summed up in Table 3.

Our analyses did not suggest any sign of selective disappearance, as the age at last observation was never retained in the selected models, except for alpha1-globulins (β ± standard-error (SE) = −0.02 ± 0.01; Table 1). Some immune trait values were impacted by the delay between capture and sampling (neutrophils: β ± SE = 0.04.10^−1^ ± 0.06.10^−2^; basophils: β ± SE = 0.08.10^−3^ ± 0.03.10^−3^; eosinophils: β ± SE = −0.04.10^−2^ ± 0.05.10^−3^; HL: β ± SE = 0.08.10^−2^ ± 0.03.10^−2^; lymphocytes: β ± SE = −0.01.10^−1^ ± 0.03.10^−2^; Table 1). Body mass was negatively associated to globulin concentrations mostly, suggesting that individuals of poorest condition had lowest protein resources (alpha1-globulins: β ± SE = −0.04 ± 0.01; beta-globulins: β ± SE = −0.10 ± 0.03; gamma-globulins: β ± SE = −0.19 ± 0.06; Table 1). Only alpha1-globulins showed temporal variation throughout the capture season, with a trend to decrease across the season (β ± SE = −0.03.10^−1^ ± 0.01.10^−1^; Table 1).

The relationship between juvenile FGMs and immunosenescence rates was evaluated on 249-332 observations from 136-159 individuals, depending on the investigated trait (see Table S4). The slope of the lymphocyte count after the onset of senescence (*i.e.* 4 years of age) decreased as juvenile FGMs increased (β_FGM_ ± SE = 0.19 ± 0.11; β_age.2_ ± SE = 1.77 ± 0.62; β_FGM:age.2_ ± SE = −0.24 ± 0.10; Figure 2, Table S4). For all other traits, we failed to detect a link between juvenile FGMs and immunosenescence rates (Table S4). Indeed, models including either the additive or interacting effect of juvenile FGMs with age did not improve the model fit in any case (Table S4).

**Figure 2.**
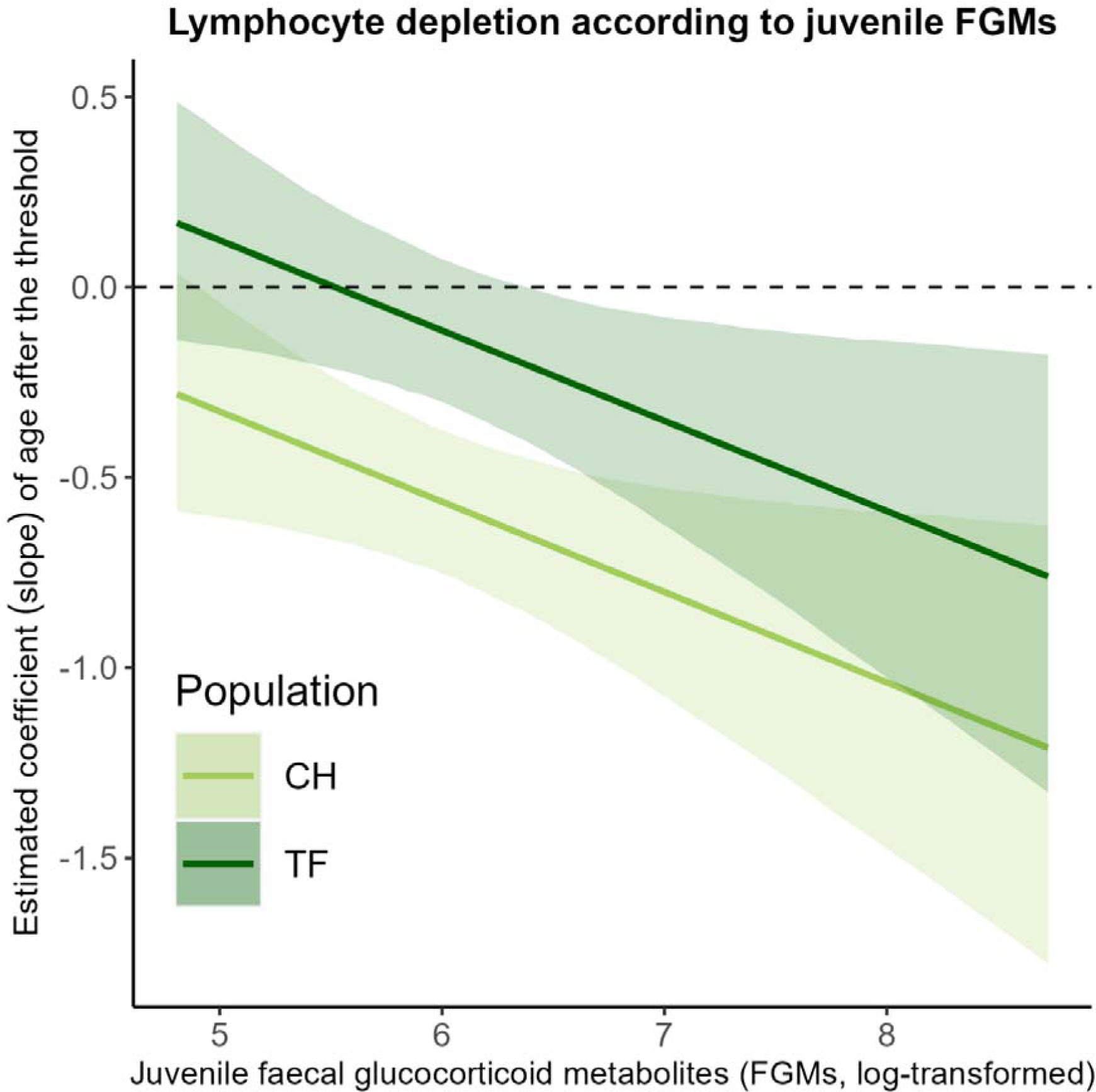
Predicted slope of lymphocyte concentration as a function of age after the threshold (*i.e.* 4 years of age, Figure 1G) according to faecal glucocorticoid metabolites (FGMs, log-transformed) measured during the first year of life and the population (CH: Chizé; TF: Trois-Fontaines). Lines are the model predictions and shaded areas are 95% CIs.

### 3.2. Parasite abundance

Age-specific patterns of parasite abundance were determined based on 915-976 observations from 449-469 individuals, depending on the parasite investigated. The abundance of GI strongyles increased with age similarly in both sexes and in both populations until 10 years of age. Afterwards, male parasitic abundance increased even more. GI strongyles abundance was consistently higher in males than in females (Table 2, Figure 3A). At CH, the low-quality habitat, the abundance of *Trichuris* sp. increased in both sexes, although more strongly in males than in females up to 10 years of age, after what parasitic abundance tended to decrease in females and to increase even more in males until the end of life. At TF, parasitic abundance in males increased up to 10 years of age and accelerated afterwards, whereas it remained constant in females up to that age and increased slightly less than for males afterwards. *Trichuris* sp. abundance was always higher in males compared to females, and in CH compared to TF (Table 2, Figure 3B). Protostrongylid abundance was higher in males than in females and increased with age in both sexes, although faster in males compared to females. Parasitic abundance increased from *ca.* 5 and 6 years of age in males and females, respectively, whereas it increased from *ca.* 2 years of age in both sexes for individuals born during years of good quality, although faster in males compared to females. Protostrongylid abundance was lower at CH than at TF, and was higher and increased more rapidly for individuals born during years of good quality compared to those born during years of poor quality (Table 2, Figure 3C). We did not detect any change in abundance of coccidia in either population or sex (Table 2, Figure 3D). Onsets and rates of change in parasite abundance with age are summed up in Table 3.

**Figure 3.**
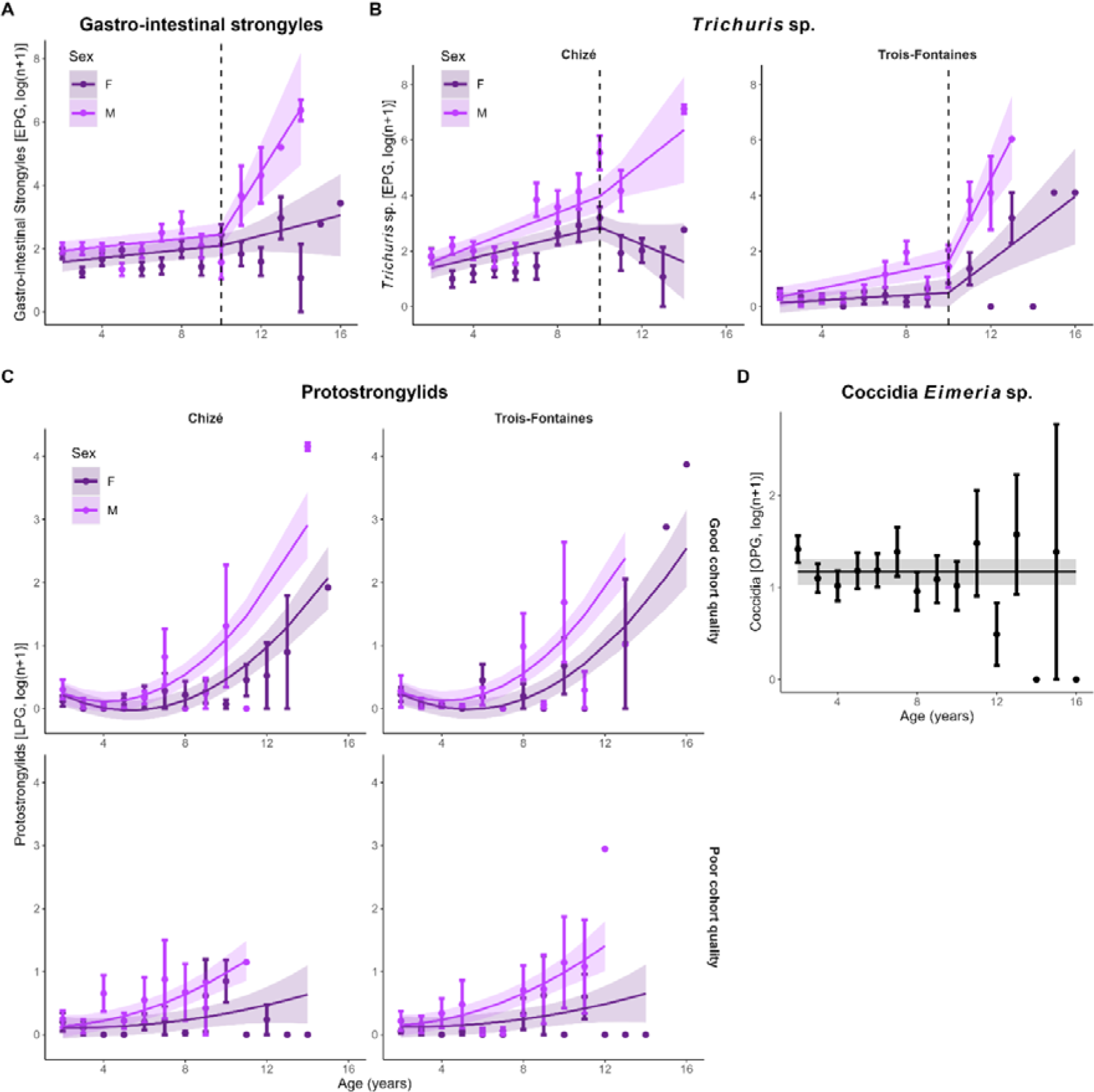
Parasite abundance age trajectories in roe deer. Lines are retained model predictions for each parasitic trait (excluding retained confounding variables for graphical representation) and shaded areas are 95% CIs. EPG: eggs per gram, LPG: larvae per gram, OPG: oocysts per gram. Points are age-specific average trait values ± standard errors. Dots without error bars correspond to unique observations.

**Table 2.**
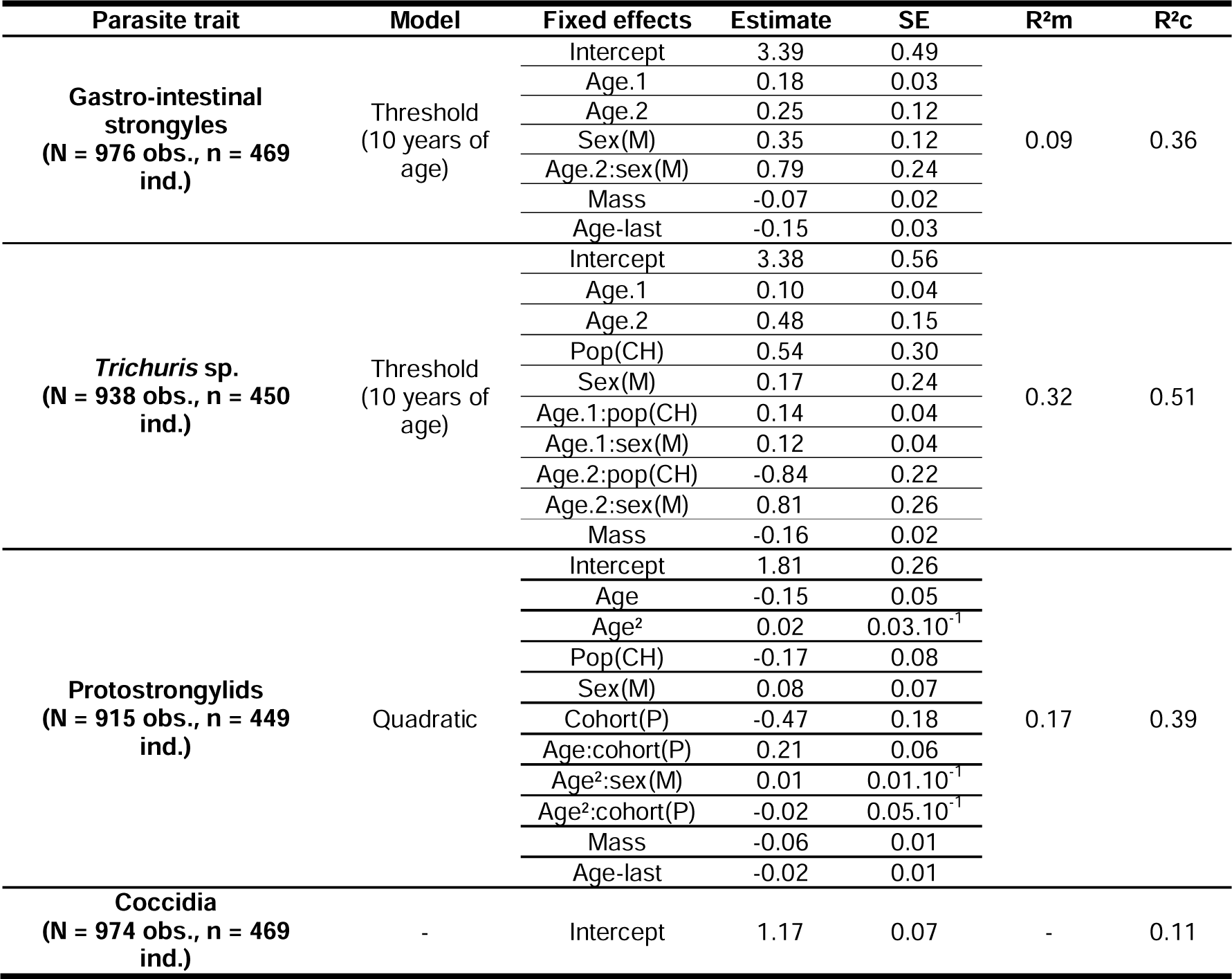
Selected linear mixed effect models for the 4 parasitic traits according to age (linear, quadratic, threshold), population (‘Pop’, reference level is Trois-Fontaines; CH: Chizé), sex (‘Sex’, reference level is females; M: males), and cohort quality (‘Cohort’, reference level is ‘good cohort quality’; P: poor cohort quality). Models were selected based on the Akaike Information Criteria corrected for small sample sizes (AICc) as described in section 2.5.2. AICc values of retained models can be found in Tables S3. Models accounted for the following potential confounding effects: body mass (‘Mass’), age at last observation (‘Age-last’) and Julian day of capture (‘Julian’). All models included the random effects of the individual identity and the year of capture nested within the population. SE: standard error, R²m – R²c: marginal – conditional model variance, respectively.

All parasite abundances, except for coccidia, were negatively related to individual body mass, with lighter individuals being more parasitised than heavier ones (GI strongyles: β ± SE = −0.07 ± 0.02; *Trichuris* sp.: β ± SE = −0.16 ± 0.02; Protostrongylids: β ± SE = −0.06 ± 0.01; Table 2). We detected signs of selective disappearance for GI strongyles (β ± SE = −0.15 ± 0.03; Table 2) and protostrongylids (β ± SE = −0.02 ± 0.01; Table 2).

The effect of juvenile FGMs on retained trajectories was evaluated on 252-274 observations from 129-140 individuals, depending on the investigated parasite (see Table S4). For protostrongylids we retained the additive effect of juvenile FGMs on adult abundance interacting with the cohort quality (β_FGM_ ± SE = 0.04 ± 0.12, β_Cohort quality(Poor)_ ± SE = −3.24 ± 1.21; β_FGM:Cohort_quality(Poor)_ ± SE = 0.50 ± 0.18; Table S4, Figure 4). This suggests that among individuals born during years of poor quality, those with higher juvenile FGMs display higher protostrongylid loads as adult, whereas no particular trend was detected for individuals born during years of good quality (Figure 4). Note that when removing the high residual value on the top right of Figure 3B, only a positive additive effect of juvenile FGMs on adult abundance was retained (β ± SE = 0.15 ± 0.08). We found no evidence for a link between juvenile FGMs and GI nematodes or protozoan abundance trajectories (Table S4).

**Figure 4.**
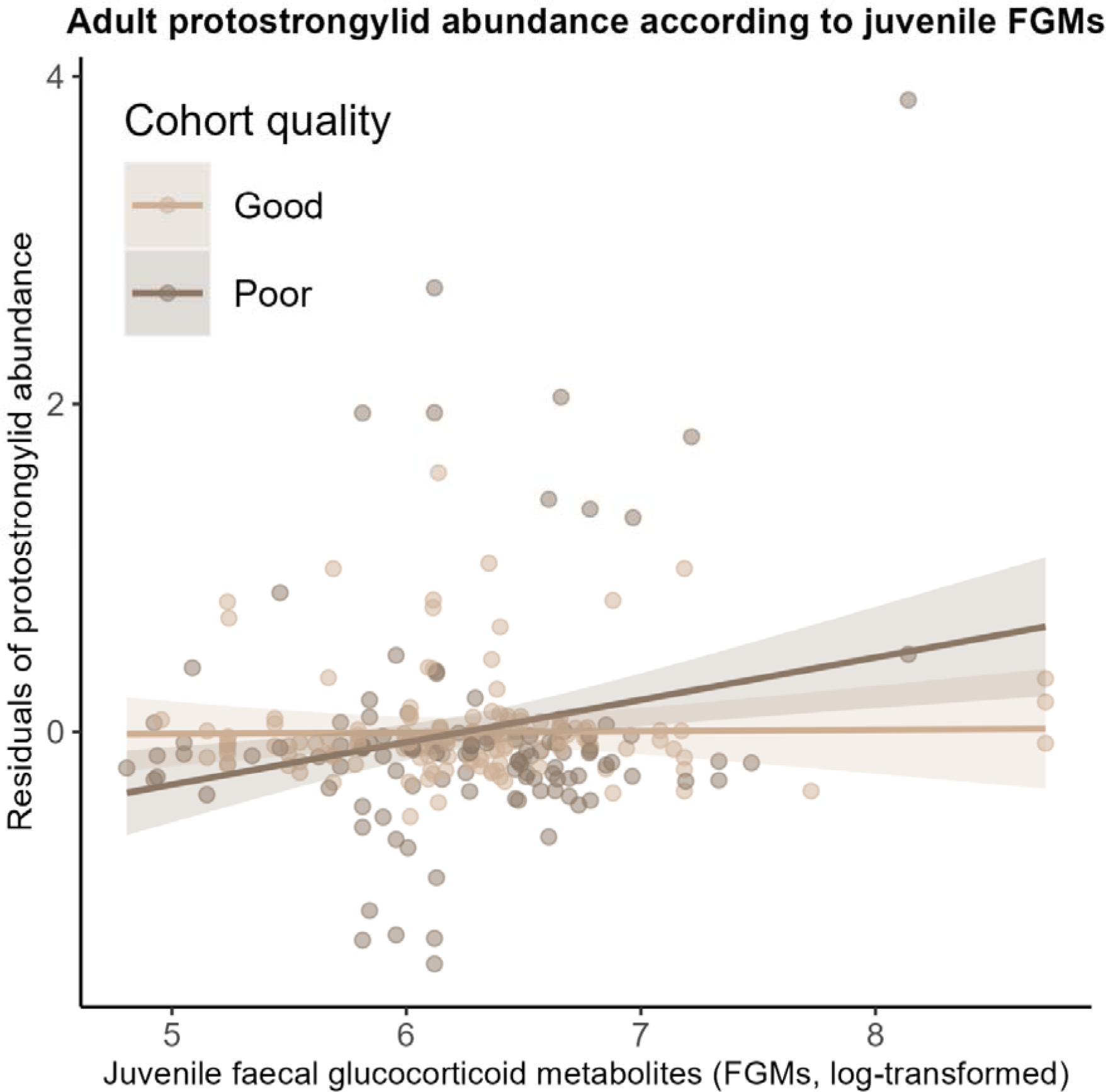
Predicted roe deer protostrongylid abundance accounting for age, population, sex, cohort, body mass and age at last observation (residuals of the retained parasite trajectory, Figure 2C) according to FGMs (log-transformed) measured during the first year of life and cohort quality. Note that when removing the particularly high parasite abundance value (*i.e.* top right corner), no cohort effect was retained. Lines are the model predictions and shaded areas are 95% CIs.

## 4. Discussion

There is a gap in the literature on the putative relationship between early-life GC levels and immunosenescence in mammals. Studies that demonstrated negative relationships between GCs and immunosenescence focused mostly on humans (Bauer 2005, 2008, Garrido et al. 2022), and only few studies, all conducted on birds, specifically assessed the effect of stress exposure or GC levels during the juvenile period on late-life performance, evidencing negative relationships (Spencer et al. 2009, Monaghan et al. 2011, Herborn et al. 2014, Monaghan and Haussmann 2015, Casagrande et al. 2020, but see also Spencer and Verhulst 2008). Here, we explored whether GCs, measured as FGMs during the juvenile period, were linked to patterns of immunosenescence in wild roe deer. We used a remarkably large dataset and found that, overall, patterns of immunosenescence supported those previously reported by Cheynel et al. (2017) in roe deer, although few differences and novelties arose (*e.g.* sex and population effects, different shape trajectories). Most importantly, increased juvenile FGM levels were associated to accelerated depletion of the lymphocyte pool after 4 years of age, but were not related to the senescence patterns of other immune traits measured. Also, FGM levels in juveniles were positively associated, although weakly, with the abundance of pulmonary parasites (*i.e.* protostrongylids), measured during adulthood for individuals born during years of poor quality, but without modulating the ageing trajectory of these parasite loads.

Lymphocyte count was higher in the good-quality habitat (*i.e.* at TF) compared to the low-quality habitat (*i.e.* at CH), and higher in females compared to males. For both sexes and populations, lymphocyte count decreased linearly from to 2 to 4 years of age, after what count decreased at CH and increased at TF. A decrease was expected because of the thymic involution occurring with advancing age. It results in a decreased production of naïve B and T lymphocyte and changes in the lymphocyte pool population (Hakim and Gress 2007). By contrast, the increase observed at TF rather suggests that the expected decline in lymphocyte count can be dampened or prevented by the more favourable environmental conditions encountered at TF (McDade et al. 2016). Gamma-globulin concentration increased with age in both populations and was higher at CH than at TF, while lymphocyte count was lower at CH than at TF. Thus, B lymphocytes from individuals of CH could produce more antibodies relative to their number compared to TF, potentially resulting in a stronger auto-immunity at CH than at TF. Indeed, immunosenescence has been found to be accompanied by a decrease in the production of high-affinity antibodies (Frasca et al. 2011) and by an increased production of auto-antibodies, associated to autoimmune diseases in some cases (Hakim and Gress 2007, Kovaiou et al. 2007, Lleo et al. 2010, Bachi et al. 2013). This suggests a dysregulation rather than a mere decline of the adaptive immunity with increasing age. Nevertheless, we do not have sufficient information about the lymphocyte pool composition (*i.e.* relative proportions of NK cells, B and T lymphocytes and proportions of naïve *v.* mature lymphocytes) to draw strong conclusions. Moreover, additional characterisation of the immunoglobulins (*e.g.* isotypes, affinity, glycolisation) would be necessary to evaluate their potential efficiency against certain parasites or ability to induce an autoimmune response. Still, altogether, our results suggest that the adaptive immunity functioning could be altered between populations, even if in different ways, in line with a decline in adaptive immunity with increasing age (Peters et al. 2019).

Age-related variation in lymphocyte count beyond 4 years of age was also modulated by FGM exposure during the juvenile stage. As juvenile FGMs increased, lymphocyte count decreased at a faster rate with age. Lymphocytes are especially sensitive to chronically elevated GCs, which promote thymocyte (*i.e.* immature lymphocyte within the thymus) apoptosis and thymic involution (Gruber et al. 1994, Gomez-Sanchez 2009, Taves et al. 2017), whereas GCs below stress-induced levels seem to rather enhance survival of lymphocytes and to delay thymic involution (Taves et al. 2017). Thymus is the primary lymphoid organ responsible for the maturation and selection of T cells (Palmer 2013), suggesting that the age-related depletion of the lymphocyte pool can be accelerated by GCs. GCs are primarily produced by the adrenal glands but are also locally produced, notably by the thymus. Chronic elevation of adrenal GCs, such as during adverse environmental conditions, can result in an increased thymic GC production, amplifying the adverse effect of GCs on thymus functioning (Taves et al. 2017). This increased thymic sensitivity to elevated GCs can partly explain the observed effect of juvenile GCs on lymphocytes but not on other immune traits. Investigating how adrenal and thymic GC productions covary in these populations could help to better understand how environmental conditions relate to changes in lymphocyte count.

FGMs during the juvenile period were weakly but positively related to the abundance of lung parasites during adulthood in a cohort-specific manner: juveniles with higher FGM levels exhibited a higher abundance of protostrongylids during adulthood than juveniles with lower FGM levels, only for individuals born during years of poor quality. Protostrongylids have a long period of patency (> 2 years) and do not enable the final host to develop acquired immunity (Adem 2016). In the absence of adaptive immunity against these parasites and because young individuals are expected to rely more heavily on innate rather than adaptive immunity (McDade et al. 2016), high juvenile FGMs could disrupt early innate immunity (Carbillet et al. 2023a), resulting in individuals more heavily infected during adulthood. Our results suggest that this is even more the case for individuals born during years of poor quality for which innate immunity might be more sensitive to glucocorticoids (Carbillet et al. 2023a), although the cohort effect appeared to be mainly driven by one heavily parasitised individual from a poor cohort.

Protostrongylids are weakly pathogenic parasites, except when individuals are heavily infected (Adem 2016). We found evidence of selective disappearance (β = −0.02, se = 0.01, Table 2), with surviving individuals at advanced age being the least parasitised. This can explain that individuals born during years of poor quality displayed lower parasitic loads compared to others. Individuals from poor quality years might not survive heavy parasite loads, whereas individuals from good cohorts may better cope with the costs of being heavily parasitised. Similarly, selective disappearance occurred for GI strongyles (β = −0.13, se = 0.03, Table 2) and was stronger than for protostrongylids. Thus, individuals heavily parasitised by GI strongyles are likely under-represented in our dataset, masking a potential effect of juvenile GCs on adult abundance. GI strongyle species are diverse in both populations and some of these species are highly pathogenic (*e.g. Bunostomum trigonocephalum*, *Haemonchus contortus*; Beaumelle et al. 2021). Hence, the selective pressure of parasites on individual capacity to maintain efficient immunity, regardless of GC levels, could be higher for GI strongyles than other parasites. Interestingly, these highly pathogenic parasites, generally associated to domestic ruminants, were only found at TF (Beaumelle et al. 2021). Roe deer contamination can have occurred in TF as domestic sheep have been bred in the surrounding of the study site (Beaumelle et al. 2021). Contamination might also occur in CH but these parasites could only establish in TF where individuals are of better quality and better able to survive infection.

Overall, except for lymphocyte count, we observed a lack of relationship between juvenile FGMs and immunosenescence, and this could be due to a certain number of limitations related to GCs. First, the stress response does not only involve the HPA axis and GCs but also other hormones, cytokines and neurotransmitters (Sapolsky et al. 2000). Similarly, GCs are not the sole contributors to allostasis, and estimating allostatic load should ideally be done using several markers to summarise the level of physiological activity (Seeman et al. 1997, 2001, Romero et al. 2009, Juster et al. 2010). Secondly, it is admitted that acute GCs elevations can stimulate immune functions, whereas high baseline GC levels or chronic GC elevations often result in immunosuppression (Martin 2009, Dhabhar 2014). However, chronic/long-term GC elevations have also been reported to decrease the responsiveness of macrophages to GCs, resulting in an up-regulation of immune activity as expected for individuals living at risk of disease or infection threats to ensure competent immune system at all times (Martin et al. 2005, Martin 2009). Thirdly, the consequences of GCs on certain traits, such as immune ones, could be restricted to immediate or short-term consequences (*i.e.* days-weeks), but could be lifelong on others, such as telomere dynamics, as evidenced in the same roe deer populations (Lemaître et al. 2021). Also, the timing of FGM measurement, at 8 months of age, could be too late to assess carry-over consequences of GCs on physiology. Such consequences could be restricted to very early-life stages during the HPA axis maturation (Novais et al. 2017) while roe deer appear to produce relatively mature offspring regarding neuroendocrine functions (Bonnot et al. 2018). Finally, it is important to note that juvenile FGMs are measured on individuals that have survived up to that age, following a strong viability selection (Garratt et al. 2015).

A recent study on the same two roe deer populations has shown that baseline GC levels were negatively associated with individual body mass, but GCs measured during the first year of life showed no delayed relationship with adult body mass (Lalande et al. 2023). Similarly, we were not able to detect any global carry-over effects of GCs on roe deer immunity and immunosenescence except for lymphocytes, while, over a short period of time (*i.e.* 8 weeks), baseline GC levels have been shown to positively covary over time with innate and adaptive immunity, but not inflammatory response, in this species (Carbillet et al. 2022). The time window considered can possibly explain the overall observed absence of relationship, as many events can happen between the juvenile and senescent stage. Among these events, most juvenile natal dispersal occurs about three months after juvenile FGMs are measured (Hewison et al. 2021). Dispersal can influence both current (Maag et al. 2019) and future FGM levels depending on the local environmental conditions dispersers settle in, especially in the heterogeneous forest of CH.

By considering a larger dataset and more confounding factors, we found patterns of immunosenescence showing the same trends as those previously found by Cheynel et al. (2017), but with slightly different age trajectories for some traits (beta-globulins, haptoglobins, gamma-globulins, lymphocytes GI strongyles, *Trichuris* sp. and protostrongylids). However, some differences, likely due to an increase in sample size (by 66-95% depending on the trait), deserve to be noted. First, monocytes that previously showed no changes with age displayed senescence here (Table 3). Secondly, while Cheynel et al. (2017) showed population-specific trajectories for six traits, it was the case only for lymphocyte count and *Trichuris* sp. abundance here, so that population did not seem to be the main modulator of senescence patterns anymore. Thirdly, we found an effect of natal environmental conditions on protostrongylid trajectories but not on the value or senescence of other immune and parasite traits. Yet, environmental conditions at the time of birth likely affect resource availability and quality, which can interfere with lifetime trajectories of life-history traits (Nussey et al. 2007, Cooper and Kruuk 2018), and notably changes in the immune system with age (Lochmiller and Deerenberg 2000, McDade 2005, Rauw 2012, McDade et al. 2016).

**Table 3.** Summary of evidence for immunosenescence and change in parasite abundances with age in roe deer. 12 immune traits and four parasites were investigated. Estimations of onsets and rates of changes with age are provided based on retained model predictions. F: females; M: males, CH: Chizé, TF: Trois-Fontaines, EPG: eggs per gram of faeces, OPG: oocysts per gram of faeces. For quadratic trajectories, onsets were estimated as the age for which the derivative was null. Rates were estimated using the derivative of the function for the onset and for the last age available to represent the range of change with age from the onset to the end of life. When the onset was <2, rates were estimated at 2 years of age.

| Trait | Immunosenescence<br>/<br>Change in parasite<br>abundance with age | Onset (years) | Rate |
| --- | --- | --- | --- |
| **INNATE IMMUNITY** |  |  |  |
| Neutrophil | Yes | <2 | +28–224 cells/year |
| Monocyte | Yes | 2 | -20 cells/year |
| Basophil | No | - | - |
| Eosinophil | No | - | - |
| Haemagglutination | No | - | - |
| Haemolysis | No | - | - |
| **INFLAMMATORY RESPONSE** |  |  |  |
| Alpha1-globulin | Yes | <2 | F: +0.02–0.13 mg.mL<sup>-1</sup>/year;<br>M: +0–0.04 mg.mL<sup>-1</sup>/year |
| Alpha2-globulin | No | - | - |
| Beta-globulin | Yes | 2 | +0.23 mg.mL<sup>-1</sup>/year |
| Haptoglobin | Yes | <2 | +0.02–0.13 mg.mL<sup>-1</sup>/year |
| **ADAPTIVE IMMUNITY** |  |  |  |
| Gamma-globulin | Yes | 2 | +0.42 mg.mL<sup>-1</sup>/year |
| Lymphocyte | Yes | 4 | CH: -20 cells/year;<br>TF: +60 cells/year |
| **PARASITES** |  |  |  |
| Gastro-intestinal strongyles | Yes | 10 | F: +0.28 EPG/year<br>M: +1.86 EPG/year |
| *Trichuris* sp. | Yes | 10 | F-CH: -0.30 EPG/year;<br>F-TF: +0.61 EPG/year;<br>M-CH: +0.58 EPG/year;<br>M-TF: +2.63 EPG/year |
| Protostrongylids | Yes | Good cohort:<br>F = 4.2; M =<br>3.2<br>Poor cohort:<br><2 | Good cohort-F: +0.00–0.53<br>LPG/year;<br>Good cohort-M: +0.00–0.60<br>LPG/year;<br>Poor cohort-F: +0.06–0.07<br>LPG/year;<br>Poor cohort-M: +0.09–0.23<br>LPG/year |
| Coccidia | No | - | - |

Immune traits displayed no sex-specific age trajectories, except for alpha1-globulins. In females, alpha1-globulins displayed lower concentration at young age but increased later in life until catching up and going beyond male values, although this increase was weak due to marked age-related variation. The previously documented sex-specific trajectory for neutrophils is no longer observed in the current study, while it remained similar for *Trichuris* sp. and protostrongylids, and even extended to GI strongyles, so that sex, as previously found, is not a modulator of immunosenescence but remains the main modulator of parasite changes with age. According to the immunocompetence handicap hypothesis, males should display a weaker immunity due to allocation trade-offs between immunity and reproduction (Zuk 1990) and the detrimental effects of male sex hormones on immunity (Foo et al. 2017). However, empirical evidence for a stronger female immunocompetence is mixed, and a recent meta-analysis actually showed no detectable bias towards females (Kelly et al. 2018). Similarly, although sex-specific patterns of senescence are expected from sex-specific life-histories (Brooks and Garratt 2017), a meta-analysis showed no bias towards an earlier or accelerated immunosenescence in males compared to females (Peters et al. 2019). In line with this, we showed that, throughout life, immune trait values were higher (alpha1-globulins, beta-globulins, haptoglobins), similar (neutrophils, monocytes, basophils, HA and HL scores, gamma-globulins and coccidia), or lower (eosinophils, alpha2-globulins, lymphocytes) in males compared to females in this weakly sexually dimorphic species (Andersen et al. 1998). However, except for coccidia, parasite load was constantly greater in males than in females, and even increased more rapidly with age for males (*Trichuris* sp. and protostrongylids). This is unlikely due to sex-specific behaviours that could result in different parasite exposure between males and females, as both sexes do not segregate spatially throughout the year (Bonenfant et al. 2007). This rather suggests that although immune trait values do not strongly differ between sexes, resistance to parasite could actually be stronger in females than in males, as it has previously been suggested in other species (Poulin 1996, Zuk and McKean 1996, Tidière et al. 2020).

Finally, immunosenescence patterns for innate and inflammatory traits are consistent with the trajectories described in the literature on wildlife, humans or laboratory organisms (Peters et al. 2019), with innate traits showing no particular ageing directions, and inflammatory markers increasing with advancing age (Table 3). Also, neutrophil count was higher at the good-quality environment encountered at TF than at CH, suggesting a poorer phagocytic capacity in the latter population, and potentially a poorer immunocompetence, hence supporting that low resource quality and availability at CH can impact some immune traits (McDade et al. 2016). Similarly, we evidenced that parasite load was greater in CH than TF at all ages for *Trichuris* sp. and protostrongylids, but not for coccidia and GI strongyles (showing no population-specific differences), a pattern recently reported in those two populations (Bariod et al. 2023). Whether this is due to different pathogen exposure between populations or whether it is a consequence of a poorer immunocompetence in CH compared to TF remains to be assessed.

## Ethics

The research presented in the current article was done according to all institutional and/or national guidelines. For both populations (*i.e.* Trois-Fontaines and Chizé), the protocol of capture and blood sampling of roe deer is under the authority of the *Office Français de la Biodiversité* (OFB) and was approved by the Director of Food, Agriculture and Forest (Prefectoral order 2009-14 from Paris). All procedures were approved by the Ethical Committee of Lyon 1 University (project DR2014-09, June 5^th^, 2014 / MESRI #40903-2023021016306261, March 7^th^, 2023).

## Acknowledgements

This work was conducted as part of a PhD funded by the *Université de Lyon* and the *Office Français de la Biodiversité* (OFB). We are grateful to all technicians, researchers and volunteers participating in the collect of data on both sites (Trois-Fontaines and Chizé). The Trois-Fontaines population is part of the long-term Studies in Ecology and Evolution (SEE-Life) program of the CNRS.

## Author contributions

**Conceptualisation:** LL, EGF and PV

**Field sampling:** GB, JC, LC, FD, JMG, RG, EGF, LL, JFL, MP, CP, BR and PV

**Laboratory:** GB, JC, LC, HF, EGF, LL, RP, CP, BR and PV

**Data curation:** GB, EGF and LL

**Statistical analysis:** LL

**Funding acquisition:** EGF, JFL and PV

**Methodology:** EGF, LL and PV

**Project administration:** JMG, EGF, JFL, MP and PV

**Supervision:** EGF and PV

**Writing – original draft:** LL

**Writing – review and editing:** GB, JC, LC, JMG, EGF, LL, JFL, RP, CP, BR and PV

