## Supplementary Information for "Early-life glucocorticoids accelerate lymphocyte count senescence in roe deer"

**Table S1.** Estimation of the number of roe deer (> 1 year of age; N) before population management (hunting) in Chizé and Trois-Fontaines across the capture period considered in the present analysis (2010-2022). λ: population growth rate.

| Year of capture | Chizé | | Trois-Fontaines | |
|---|---|---|---|---|
| | N [min, max] | λ | N [min, max] | λ |
| **2010** | 473 [386, 597] | 1.57 | 164 [141, 199] | 0.94 |
| **2011** | 386 [331, 463] | 0.82 | 197 [169, 239] | 1.20 |
| **2012** | 431 [343, 558] | 1.12 | 269 [213, 351] | 1.37 |
| **2013** | 330 [278, 406] | 0.77 | 232 [196, 284] | 0.86 |
| **2014** | 339 [287, 413] | 1.03 | 239 [202, 296] | 1.03 |
| **2015** | 327 [284, 392] | 0.96 | 143 [121, 176] | 0.60 |
| **2016** | 372 [312, 413] | 1.14 | 180 [156, 217] | 1.26 |
| **2017** | 348 [298, 421] | 0.94 | 172 [149, 207] | 0.96 |
| **2018** | 459 [388, 561] | 1.32 | 222 [188, 270] | 1.29 |
| **2019** | 377 [301, 488] | 0.82 | 201 [170, 246] | 0.91 |
| **2020** | 317 [263, 398] | 0.84 | 235 [186, 307] | 1.17 |
| **2021** | 394 [242, 672] | 1.24 | 256 [180, 375] | 1.09 |
| **2022** | 299 [255, 381] | 0.76 | 311 [248, 403] | 1.21 |

**Table S2.** Linear mixed effects model (LMM) selection table for the correction of faecal glucocorticoid metabolites measured during the juvenile stage (FGMs, log-transformed, N = 162 observations on 162 individuals) according to Julian date of capture (linear and quadratic), the time delay between capture and sampling ('Delay' in minutes) and whether faecal samples were immediately frozen at -80°C or frozen within 24 hours after collection ('Freezing'). Values give the parameter coefficient and values between brackets are the standard-errors. 'I' is the Intercept, 'df' is the number of parameters, 'Log-lik' is the log-likelihood, 'Delta' is the difference of AICc between the candidate model and the model having the lowest AICc, and 'AICcw' the AICc weight of each model. Retained model is in bold. Dashed line separate models below 2 ΔAICc and models above.

| I | Julian | Julian² | Delay | Freezing(Immediate) | df | Log-lik | AICc | Delta | AICcw |
|---|---|---|---|---|---|---|---|---|---|
| **6.39 (0.05)** | | | | | **2** | **-156.25** | **316.57** | **0.00** | **0.19** |
| 6.18 (0.17) | $0.03.10^{-1}$ ($0.02.10^{-1}$) | | | | 3 | -155.41 | 316.97 | 0.40 | 0.15 |
| 6.28 (0.10) | | $0.02.10^{-3}$ ($0.02.10^{-3}$) | | | 3 | -155.52 | 317.18 | 0.62 | 0.14 |
| 6.35 (0.09) | | | | 0.06 (0.11) | 3 | -156.10 | 318.36 | 1.79 | 0.08 |
| 6.35 (0.15) | | | $0.01.10^{-2}$ ($0.05.10^{-2}$) | | 3 | -156.21 | 318.58 | 2.01 | 0.07 |
| 6.17 (0.17) | $0.03.10^{-1}$ ($0.02.10^{-1}$) | | | 0.03 (0.11) | 4 | -155.38 | 319.01 | 2.45 | 0.06 |
| 6.12 (0.33) | $0.06.10^{-1}$ (0.01) | $-0.02.10^{-3}$ ($0.09.10^{-3}$) | | | 4 | -155.38 | 319.01 | 2.45 | 0.06 |
| 6.15 (0.21) | $0.03.10^{-1}$ ($0.02.10^{-1}$) | | $0.01.10^{-2}$ ($0.05.10^{-2}$) | | 4 | -155.39 | 319.02 | 2.46 | 0.05 |
| 6.26 (0.12) | | $0.02.10^{-3}$ ($0.02.10^{-3}$) | | 0.03 (0.11) | 4 | -155.48 | 319.22 | 2.66 | 0.05 |
| 6.25 (0.17) | | $0.02.10^{-3}$ ($0.02.10^{-3}$) | $0.01.10^{-2}$ ($0.05.10^{-2}$) | | 4 | -155.50 | 319.25 | 2.69 | 0.05 |
| 6.29 (0.18) | | | $0.01.10^{-2}$ ($0.05.10^{-2}$) | 0.06 (0.11) | 4 | -156.04 | 320.34 | 3.77 | 0.03 |
| 6.14 (0.23) | $0.03.10^{-1}$ ($0.02.10^{-1}$) | | $0.01.10^{-2}$ ($0.05.10^{-2}$) | 0.03 (0.11) | 5 | -155.34 | 321.07 | 4.51 | 0.02 |
| 6.10 (0.33) | $0.06.10^{-1}$ (0.01) | $-0.02.10^{-3}$ ($0.09.10^{-3}$) | | 0.03 (0.11) | 5 | -155.35 | 321.08 | 4.51 | 0.02 |
| 6.07 (0.37) | $0.06.10^{-1}$ (0.01) | $-0.02.10^{-3}$ ($0.09.10^{-3}$) | $0.01.10^{-2}$ ($0.05.10^{-2}$) | | 5 | -155.35 | 321.08 | 4.51 | 0.02 |
| 6.23 (0.19) | | $0.02.10^{-3}$ ($0.02.10^{-3}$) | $0.01.10^{-2}$ ($0.05.10^{-2}$) | 0.03 (0.11) | 5 | -155.45 | 321.29 | 4.73 | 0.02 |
| 6.05 (0.38) | $0.06.10^{-1}$ (0.01) | $-0.03.10^{-3}$ ($0.09.10^{-3}$) | $0.02.10^{-2}$ ($0.06.10^{-2}$) | 0.03 (0.11) | 6 | -155.30 | 323.14 | 6.58 | 0.01 |

**Table S3.** Model selection based on AICc among candidate linear mixed models (LMM) testing the relationship between immune and parasite traits and different age functions (*i.e.* no age variations, linear, quadratic, threshold age functions) in Trois-Fontaines and Chizé. Retained models are in bold. Thr: threshold age (in years), k: number of model parameters, ΔAICc: AICc difference between the candidate model and the model with the lowest AICc, AICcw: AICc weight to measure the relative likelihood of each model to be the best model among the set of fitted models. All full models included the different age functions mentioned, the additive effects of the population ('pop'), the sex ('sex'), the cohort quality ('cohort') and the two-way interactions between these effects and age terms. Full models also included confounding variables as defined in Material and Methods (*i.e.* 'mass', 'age-last', 'delay', 'julian'). For each age function, the table displays the most parsimonious model within 2 ΔAICc as described in the Material and Methods section.

| Age function | Neutrophil | | | | | |
|---|---|---|---|---|---|---|
| | Variables | Thr | k | AICc | ΔAICc | AICcw |
| Constant | Pop + age-last + delay | - | 7 | 4306.95 | 9.58 | 0.00 |
| Linear | Age + pop + cohort + delay | - | 8 | 4298.06 | 0.69 | 0.28 |
| Quadratic | **Age² + pop + delay** | **-** | **7** | **4297.36** | **0.00** | **0.40** |
| Post-threshold slope | Age.2 + pop + delay | 4 | 8 | 4299.15 | 1.78 | 0.16 |
| Pre-threshold slope | Age.1 + pop + cohort + delay | 10 | 9 | 4302.88 | 5.52 | 0.03 |
| Threshold, 2 slopes | Age.1 + age.2 + pop + cohort + delay | 10 | 10 | 4299.76 | 2.39 | 0.12 |
| | Monocyte | | | | | |
| | Variables | Thr | k | AICc | ΔAICc | AICcw |
| Constant | - | - | 4 | 158.73 | 7.07 | 0.02 |
| Linear | Age | - | 5 | 155.23 | 3.57 | 0.09 |
| Quadratic | Age + age² | - | 6 | 154.23 | 2.57 | 0.15 |
| Post-threshold slope | Age.2 | 6 | 6 | 161.63 | 9.97 | 0.00 |
| Pre-threshold slope | **Age.1** | **5** | **6** | **151.66** | **0.00** | **0.53** |
| Threshold, 2 slopes | Age.1 + age.2 | 5 | 7 | 153.42 | 1.76 | 0.22 |
| | Basophil | | | | | |
| | Variables | Thr | k | AICc | ΔAICc | AICcw |
| Constant | **Delay** | **-** | **5** | **-2081.26** | **1.07** | **0.18** |
| Linear | Age + delay | - | 6 | -2082.33 | 0.00 | 0.30 |
| Quadratic | Age² + delay | - | 6 | -2082.27 | 0.07 | 0.29 |
| Post-threshold slope | Age.2 + delay | 7 | 7 | -2080.19 | 2.14 | 0.10 |
| Pre-threshold slope | Age.1 + delay | 4 | 7 | -2079.60 | 2.73 | 0.08 |
| Threshold, 2 slopes | Age.1 + age.2 + delay | 4 | 8 | -2078.49 | 3.85 | 0.04 |
| | Eosinophil | | | | | |

| | Variables | Thr | k | AICc | ΔAICc | AICcw |
|---|---|---|---|---|---|---|
| Constant | **Sex + delay** | **-** | **6** | **-909.68** | **0.40** | **0.27** |
| Linear | Age + dsex + delay | - | 7 | -910.08 | 0.00 | 0.33 |
| Quadratic | $Age^2$ + sex + delay | - | 7 | -909.28 | 0.79 | 0.22 |
| Post-threshold slope | Age.2 + sex + delay | 9 | 8 | -905.66 | 4.42 | 0.04 |
| Pre-threshold slope | Age.1 + sex + delay | 3 | 8 | -906.24 | 3.83 | 0.05 |
| Threshold, 2 slopes | Age.1 + age.2 + sex + delay | 9 | 9 | -907.32 | 2.75 | 0.08 |

Haemagglutination

| | Variables | Thr | k | AICc | ΔAICc | AICcw |
|---|---|---|---|---|---|---|
| Constant | **-** | **-** | **4** | **3883.18** | **0.00** | **0.30** |
| Linear | Age | - | 5 | 3885.20 | 2.02 | 0.11 |
| Quadratic | $Age^2$ | - | 5 | 3885.18 | 2.00 | 0.11 |
| Post-threshold slope | Age.2 | 9 | 6 | 3886.93 | 3.74 | 0.05 |
| Pre-threshold slope | Age.1 + cohort + age.1:cohort | 3 | 8 | 3883.70 | 0.52 | 0.23 |
| Threshold, 2 slopes | Age.1 + age.2 + cohort + age.1:cohort + age.2:cohort | 5 | 10 | 3884.02 | 0.84 | 0.20 |

Haemolysis

| | Variables | Thr | k | AICc | ΔAICc | AICcw |
|---|---|---|---|---|---|---|
| Constant | **Serum_color + delay** | **-** | **6** | **3353.09** | **0.00** | **0.30** |
| Linear | Age + serum_color + delay | - | 7 | 3353.84 | 0.75 | 0.20 |
| Quadratic | $Age^2$ + serum_color + delay | - | 7 | 3353.94 | 0.85 | 0.19 |
| Post-threshold slope | Age.2 + serum_color + delay | 5 | 8 | 3355.87 | 2.78 | 0.07 |
| Pre-threshold slope | Age.1 + serum_color + delay | 4 | 8 | 3356.30 | 3.21 | 0.06 |
| Threshold, 2 slopes | Age.1 + age.2 + cohort + age.1:cohort + age.2:cohort + serum_color + delay | 4 | 12 | 3355.28 | 2.19 | 0.10 |

Alpha1-globulin

| | Variables | Thr | k | AICc | ΔAICc | AICcw |
|---|---|---|---|---|---|---|
| Constant | Sex + cohort + mass + julian | - | 8 | 1694.76 | 22.14 | 0.00 |
| Linear | Age + sex + age:sex + age-last + mass + delay + julian | - | 11 | 1673.13 | 0.51 | 0.27 |
| Quadratic | **$Age^2$ + sex + $age^2$:sex + age-last + mass + julian** | **-** | **10** | **1672.62** | **0.00** | **0.35** |
| Post-threshold slope | Age.2 + sex + age.2:sex + age-last + mass + julian | 5 | 11 | 1673.25 | 0.63 | 0.25 |
| Pre-threshold slope | Age.1 + sex + age.1:sex + mass + delay + julian | 10 | 11 | 1679.19 | 5.67 | 0.01 |
| Threshold, 2 slopes | Age.1 + age.2 + sex + age.2:sex + age-last + mass + julian | 6 | 12 | 1674.88 | 2.26 | 0.13 |

| | Alpha2-globulin | | | | | |
|---|---|---|---|---|---|---|
| | Variables | Thr | k | AICc | ΔAICc | AICcw |
| Constant | **Sex** | **-** | **5** | **3702.95** | **0.76** | **0.23** |
| Linear | Age + sex | - | 6 | 3704.31 | 2.11 | 0.12 |
| Quadratic | Age² + sex | - | 6 | 3704.82 | 2.62 | 0.09 |
| Post-threshold slope | Age.2 + sex + age.2:sex | | 8 | 3704.21 | 2.02 | 0.12 |
| Pre-threshold slope | Age.1 + sex | 4 | 7 | 3702.19 | 0.00 | 0.33 |
| Threshold, 2 slopes | Age.1 + age.2 + sex | 6 | 8 | 3704.33 | 2.14 | 0.11 |

| | Betaglobulin | | | | | |
|---|---|---|---|---|---|---|
| | Variables | Thr | k | AICc | ΔAICc | AICcw |
| Constant | Sex + age-last + mass + julian | - | 8 | 4111.76 | 75.60 | 0.00 |
| Linear | **Age + sex + mass** | **-** | **7** | **4036.15** | **0.00** | **0.37** |
| Quadratic | Age + age² + sex + mass | - | 8 | 4036.59 | 0.44 | 0.30 |
| Post-threshold slope | Age.2 + sex + mass | 3 | 8 | 4045.43 | 9.28 | 0.00 |
| Pre-threshold slope | Age.1 + sex + mass | 10 | 8 | 4039.47 | 3.31 | 0.07 |
| Threshold, 2 slopes | Age.1 + age.2 + pop + sex + age.2:pop + mass | 9 | 11 | 4036.94 | 0.79 | 0.25 |

| | Haptoglobin | | | | | |
|---|---|---|---|---|---|---|
| | Variables | Thr | k | AICc | ΔAICc | AICcw |
| Constant | Sex + age-last | - | 6 | 3187.08 | 13.69 | 0.00 |
| Linear | Age + sex + mass | - | 7 | 3173.39 | 0.00 | 0.30 |
| Quadratic | **Age² + sex** | **-** | **6** | **3173.39** | **0.00** | **0.29** |
| Post-threshold slope | Age.2 + sex | 9 | 7 | 3175.39 | 2.01 | 0.11 |
| Pre-threshold slope | Age.1 + sex + mass | 10 | 8 | 3179.33 | 5.95 | 0.02 |
| Threshold, 2 slopes | Age.1 + age.2 + sex + mass | 9 | 9 | 3173.45 | 0.06 | 0.27 |

| | Gammaglobulin | | | | | |
|---|---|---|---|---|---|---|
| | Variables | Thr | k | AICc | ΔAICc | AICcw |
| Constant | Pop + age-last + mass | - | 7 | 5709.29 | 43.79 | 0.00 |
| Linear | **Age + pop + mass** | **-** | **7** | **5665.51** | **0.00** | **0.49** |
| Quadratic | Age² + pop + mass | - | 7 | 5667.70 | 2.20 | 0.16 |
| Post-threshold slope | Age.2 + pop + mass | 3 | 8 | 5666.88 | 1.38 | 0.25 |
| Pre-threshold slope | Age.1 + pop + mass | 10 | 8 | 5670.91 | 5.40 | 0.03 |
| Threshold, 2 slopes | Age.1 + age.2 + pop + mass | 10 | 9 | 5669.54 | 4.03 | 0.07 |

| | Lymphocyte | | | | | |
|---|---|---|---|---|---|---|
| | Variables | Thr | k | AICc | ΔAICc | AICcw |

| | Variables | Thr | k | AICc | ΔAICc | AICcw |
|---|---|---|---|---|---|---|
| Constant | Pop + mass + delay | - | 7 | 2679.25 | 19.26 | 0.00 |
| Linear | Age + pop + sex + age:pop + mass + delay | - | 10 | 2672.75 | 12.76 | 0.00 |
| Quadratic | Age + age$^2$ + pop + sex + age:pop + mass | - | 10 | 2665.20 | 5.21 | 0.07 |
| Post-threshold slope | Age.2 + pop + age.2:pop + mass + delay | 4 | 10 | 2677.02 | 17.02 | 0.00 |
| Pre-threshold slope | Age.1 + pop + sex + age.1:pop + delay | 6 | 10 | 2671.73 | 11.73 | 0.00 |
| Threshold, 2 slopes | **Age.1 + age.2 + pop + sex + age.2:pop + delay** | **4** | **11** | **2659.99** | **0.00** | **0.93** |

Gastro-intestinal strongyles

| | Variables | Thr | k | AICc | ΔAICc | AICcw |
|---|---|---|---|---|---|---|
| Constant | Sex + mass | - | 6 | 3680.12 | 60.19 | 0.00 |
| Linear | Age + sex + age-last + mass | - | 8 | 3627.91 | 7.97 | 0.02 |
| Quadratic | Age$^2$ + sex + age-last + mass | - | 8 | 3623.97 | 4.04 | 0.11 |
| Post-threshold slope | Age.2 + sex + age-last + mass | 3 | 9 | 3626.96 | 7.03 | 0.03 |
| Pre-threshold slope | Age.1 + sex + age-last + mass | 10 | 9 | 3640.35 | 20.41 | 0.00 |
| Threshold, 2 slopes | **Age.1 + age.2 + sex + age.2:sex + age-last + mass** | **10** | **11** | **3619.93** | **0.00** | **0.85** |

*Trichuris* sp.

| | Variables | Thr | k | AICc | ΔAICc | AICcw |
|---|---|---|---|---|---|---|
| Constant | Pop + sex + age-last + mass | - | 8 | 3575.78 | 97.33 | 0.00 |
| Linear | Age + pop + sex + age:sex + mass | - | 9 | 3491.91 | 13.46 | 0.00 |
| Quadratic | Age$^2$ + pop + sex + age$^2$:sex + mass | - | 9 | 3482.40 | 3.96 | 0.11 |
| Post-threshold slope | Age.2 + pop + sex + age.2:sex + mass | 5 | 10 | 3483.11 | 4.66 | 0.08 |
| Pre-threshold slope | Age.1 + pop + sex + age.1:pop + age.1:sex + mass | 10 | 11 | 3499.89 | 21.45 | 0.00 |
| Threshold, 2 slopes | **Age.1 + age.2 + pop + sex + age.1:pop + age.1:sex + age.2:pop + age.2:sex + mass** | **10** | **14** | **3478.44** | **0.00** | **0.81** |

Protostrongylids

| | Variables | Thr | k | AICc | ΔAICc | AICcw |
|---|---|---|---|---|---|---|
| Constant | Pop + sex + age-last + mass | - | 8 | 2018.80 | 94.04 | 0.00 |
| Linear | Age + pop + sex + age:sex + mass | - | 9 | 1954.11 | 29.35 | 0.00 |
| Quadratic | **Age + age$^2$ + pop + sex + cohort + age:cohort + age$^2$:sex + age$^2$:cohort + age-last + mass** | **-** | **14** | **1924.77** | **0.00** | **0.61** |
| Post-threshold slope | Age.2 + pop + sex + age.2:sex + mass | 7 | 10 | 1943.46 | 18.70 | 0.00 |
| Pre-threshold slope | Age.1 + pop + sex + age.1:sex + mass | 10 | 10 | 1971.43 | 46.66 | 0.00 |

| | | | | | | |
|---|---|---|---|---|---|---|
| **Threshold, 2 slopes** | Age.1 + age.2 + pop + sex + cohort + age.2:sex + age.2:cohort + age-last + mass | 9 | 14 | 1925.70 | 0.93 | 0.39 |

| | Coccidia | | | | | |
|---|---|---|---|---|---|---|
| | Variables | Thr | k | AICc | ΔAICc | AICcw |
| **Constant** | **-** | **-** | **4** | **4060.78** | **0.37** | **0.25** |
| **Linear** | Age | - | 5 | 4061.63 | 1.22 | 0.16 |
| **Quadratic** | $Age^2$ | - | 5 | 4062.06 | 1.66 | 0.13 |
| **Post-threshold slope** | Age.2 | 6 | 6 | 4064.36 | 3.95 | 0.04 |
| **Pre-threshold slope** | Age.1 | 3 | 6 | 4060.41 | 0.00 | 0.30 |
| **Threshold, 2 slopes** | Age.1 + age.2 | 3 | 7 | 4062.42 | 2.01 | 0.11 |

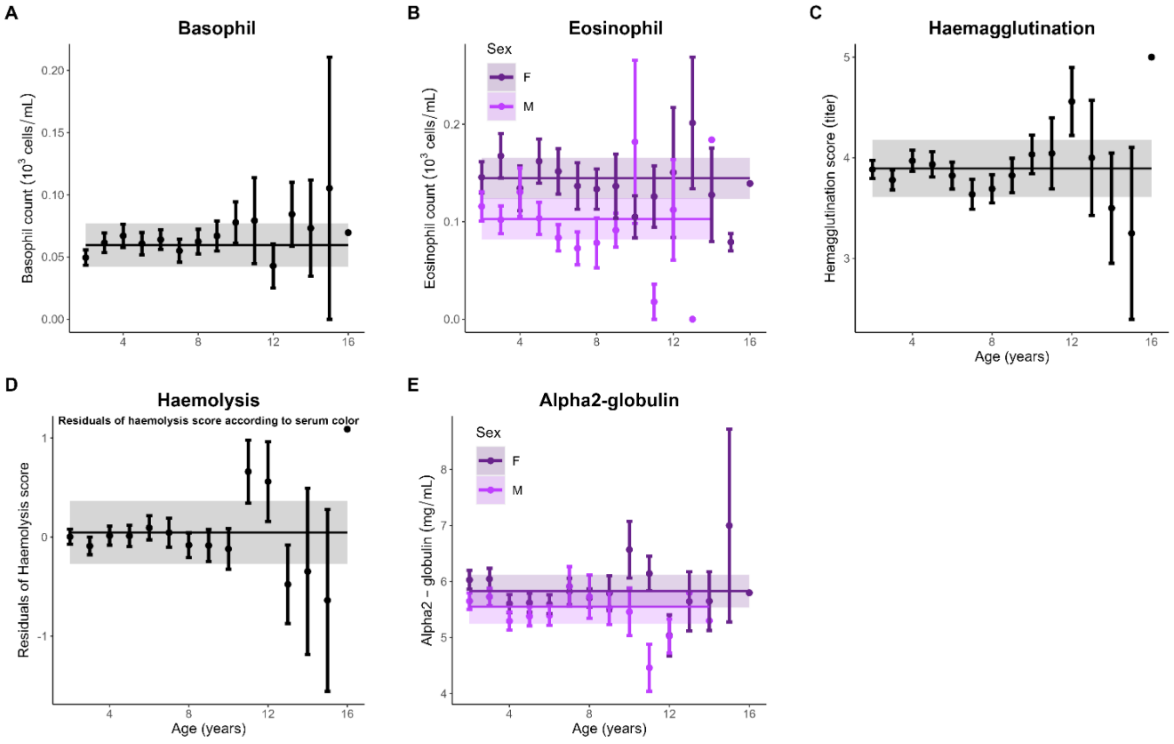


**Figure S1.** Immunosenescence patterns in roe deer. Lines are retained model predictions for each immune trait (excluding retained confounding variables for graphical representation) and shaded areas are 95% CIs. Points are age-specific average trait values ± standard errors.

**Table S4.** Linear mixed effects model (LMM) selection table for the early-late relationship between faecal glucocorticoid metabolites (FGM, log-transformed) measured during the juvenile stage and age trajectories of immune and parasite traits. Model selection was based on the ageing trajectories previously determined and included the additive effect of FGMs and the interaction between FGMs and all age terms retained in the models. 'M': Males, 'CH': population of Chizé, 'Age-last': age at last observation, 'Delay': time between capture and sampling (in minutes), 'Julian': Julian date of capture. Values give the parameter coefficient and values between brackets are the standard-errors. 'I' is the Intercept, 'df' is the number of parameters, 'Log-lik' is the log-likelihood, 'Delta' is the difference of AICc between the candidate model and the model having the lowest AICc, and 'AICcw' the AIC weight of each model. Retained model is in bold. Dashed lines separate models below 2 ΔAICc and models above.

Neutrophil (N = 267 observations, n = 142 individuals)

| I | Age² | Pop(CH) | FGM | Age²: FGM | Pop(CH): FGM | Delay | df | Log-lik | AICc | delta | AICcw |
|---|---|---|---|---|---|---|---|---|---|---|---|
| **4.91 (0.39)** | **0.01 (0.01)** | **-0.82 (0.28)** | | | | **0.00 (0.00)** | **7** | **-547.46** | **1109.34** | **0.00** | **0.38** |
| 2.87 (1.51) | 0.01 (0.01) | -0.74 (0.29) | 0.31 (0.23) | | | 0.00 (0.00) | 8 | -546.50 | 1109.57 | 0.22 | 0.34 |
| 3.28 (2.04) | 0.01 (0.01) | -1.59 (2.84) | 0.25 (0.31) | | 0.13 (0.45) | 0.00 (0.00) | 9 | -546.46 | 1111.62 | 2.28 | 0.12 |
| 3.16 (1.85) | -0.01 (0.08) | -0.74 (0.29) | 0.27 (0.28) | 0.00 (0.01) | | 0.00 (0.00) | 9 | -546.47 | 1111.63 | 2.29 | 0.12 |
| 3.54 (2.28) | -0.01 (0.08) | -1.55 (2.84) | 0.21 (0.35) | 0.00 (0.01) | 0.13 (0.45) | 0.00 (0.00) | 10 | -546.43 | 1113.71 | 4.37 | 0.04 |

Monocyte (N = 267 observations, n = 142 individuals)

| I | Age.1 | FGM | Age.1: FGM | df | Log-lik | AICc | delta | AICcw |
|---|---|---|---|---|---|---|---|---|
| **0.23 (0.05)** | **-0.04 (0.01)** | | | **5** | **-26.57** | **63.38** | **0.00** | **0.44** |
| 0.47 (0.18) | -0.04 (0.01) | -0.04 (0.03) | | 6 | -25.66 | 63.64 | 0.26 | 0.39 |
| 0.63 (0.29) | 0.05 (0.13) | -0.07 (0.05) | -0.02 (0.02) | 7 | -25.41 | 65.25 | 1.87 | 0.17 |

Basophil (N = 267 observations, n = 142 individuals)

| I | FGM | Delay | df | Log-lik | AICc | delta | AICcw |
|---|---|---|---|---|---|---|---|
| **0.11 (0.02)** | | **-0.00 (0.00)** | **5** | **260.03** | **-509.84** | **0.00** | **0.73** |
| 0.13 (0.06) | -0.00 (0.01) | -0.00 (0.00) | 6 | 260.07 | -507.82 | 2.01 | 0.27 |

Eosinophil (N = 267 observations, n = 142 individuals)

| I | Sex(M) | FGM | Sex(M): FGM | Delay | df | Log-lik | AICc | delta | AICcw |
|---|---|---|---|---|---|---|---|---|---|
| **0.31 (0.03)** | **-0.03 (0.02)** | | | **-0.00 (0.00)** | **6** | **124.15** | **-235.98** | **0.00** | **0.68** |
| 0.33 (0.11) | -0.03 (0.02) | -0.00 (0.02) | | -0.00 (0.00) | 7 | 124.17 | -233.90 | 2.08 | 0.24 |
| 0.34 (0.14) | -0.04 (0.21) | -0.00 (0.02) | 0.00 (0.03) | -0.00 (0.00) | 8 | 124.17 | -231.78 | 4.20 | 0.08 |

Hamagglutination (N = 332 observations, n = 159 individuals)

| I | FGM | df | Log-lik | AICc | delta | AICcw |
|---|---|---|---|---|---|---|
| **3.84 (0.18)** | | **4** | **-529.20** | **1066.53** | **0.00** | **0.68** |
| 4.33 (0.70) | -0.08 (0.11) | 5 | -528.93 | 1068.05 | 1.52 | 0.32 |

Haemolysis (N = 328 observations, n = 159 individuals)

| I | FGM | Delay | df | Log-lik | AICc | delta | AICcw |
|---|---|---|---|---|---|---|---|
| **0.03 (0.23)** | | **0.00 (0.00)** | **5** | **-483.90** | **977.99** | **0.00** | **0.51** |
| 0.91 (0.66) | -0.14 (0.10) | 0.00 (0.00) | 6 | -482.90 | 978.06 | 0.07 | 0.49 |

Alpha1-globulin (N = 136 observations and individuals)

| I | $Age^2$ | Sex(M) | FGM | $Age^2$: sex(M) | $Age^2$: FGM | Sex(M): FGM | Mass | Age-last | Julian | df | Log-lik | AICc | delta | AICcw |
|---|---|---|---|---|---|---|---|---|---|---|---|---|---|---|
| **3.96 (0.45)** | **0.01 (0.01)** | **0.36 (0.13)** | | **-0.01 (0.01)** | | | **-0.06 (0.02)** | **0.02 (0.02)** | **0.00 (0.00)** | **9** | **-106.90** | **233.24** | **0.00** | **0.64** |
| 3.94 (0.64) | 0.01 (0.01) | 0.36 (0.13) | 0.00 (0.07) | -0.01 (0.01) | | | -0.06 (0.02) | 0.02 (0.02) | 0.00 (0.00) | 10 | -106.90 | 235.57 | 2.33 | 0.20 |
| 4.14 (0.74) | 0.01 (0.01) | -0.10 (0.84) | -0.03 (0.10) | -0.01 (0.01) | | 0.07 (0.13) | -0.06 (0.02) | 0.02 (0.02) | 0.00 (0.00) | 11 | -106.75 | 237.63 | 4.39 | 0.07 |
| 3.80 (0.80) | 0.03 (0.06) | 0.36 (0.13) | 0.03 (0.10) | -0.01 (0.01) | -0.00 (0.01) | | -0.06 (0.02) | 0.02 (0.02) | 0.00 (0.00) | 11 | -106.86 | 237.84 | 4.61 | 0.06 |
| 4.01 (0.90) | 0.02 (0.06) | -0.07 (0.84) | -0.01 (0.12) | -0.01 (0.01) | -0.00 (0.01) | 0.07 (0.13) | -0.06 (0.02) | 0.02 (0.02) | 0.00 (0.00) | 12 | -106.72 | 239.97 | 6.74 | 0.02 |

Alpha2-globulin (N = 136 observations and individuals)

| I | Sex(M) | FGM | Sex(M): FGM | df | Log-lik | AICc | delta | AICcw |
|---|---|---|---|---|---|---|---|---|
| **5.92 (0.25)** | **-0.33 (0.24)** | | | **4** | **-247.01** | **502.33** | **0.00** | **0.60** |
| 6.49 (1.29) | -0.32 (0.24) | -0.09 (0.20) | | 5 | -246.91 | 504.28 | 1.96 | 0.23 |
| 7.89 (1.69) | -3.37 (2.40) | -0.31 (0.27) | 0.48 (0.38) | 6 | -246.10 | 504.86 | 2.53 | 0.17 |

Betaglobulin (N = 249 observations, n = 136 individuals)

| I | Age | Sex(M) | FGM | Age: FGM | Sex(M): FGM | Mass | df | Log-lik | AICc | delta | AICcw |
|---|---|---|---|---|---|---|---|---|---|---|---|
| **7.63 (0.86)** | **0.24 (0.06)** | **0.36 (0.20)** | | | | **-0.09 (0.04)** | **7** | **-462.13** | **938.72** | **0.00** | **0.53** |
| 7.76 (1.43) | 0.24 (0.06) | 0.36 (0.20) | -0.02 (0.17) | | | -0.09 (0.04) | 8 | -462.12 | 940.84 | 2.12 | 0.18 |
| 8.96 (1.66) | 0.25 (0.06) | -2.50 (2.04) | -0.22 (0.23) | | 0.45 (0.32) | -0.09 (0.04) | 9 | -461.16 | 941.07 | 2.34 | 0.16 |
| 6.94 (2.62) | 0.48 (0.64) | 0.35 (0.20) | 0.11 (0.38) | -0.04 (0.10) | | -0.09 (0.04) | 9 | -462.05 | 942.86 | 4.14 | 0.07 |
| 8.26 (2.79) | 0.44 (0.64) | -2.48 (2.04) | -0.12 (0.41) | -0.03 (0.10) | 0.45 (0.32) | -0.09 (0.04) | 10 | -461.11 | 943.14 | 4.42 | 0.06 |

Haptoglobin (N = 254 observations, n = 138 individuals)

| I | $Age^2$ | Sex(M) | FGM | $Age^2$: FGM | Sex(M): FGM | df | Log-lik | AICc | delta | AICcw |
|---|---|---|---|---|---|---|---|---|---|---|
| **0.37 (0.15)** | **0.01 (0.00)** | **0.37 (0.19)** | | | | **6** | **-425.69** | **863.71** | **0.00** | **0.61** |
| 0.39 (0.96) | 0.01 (0.00) | 0.37 (0.19) | -0.00 (0.15) | | | 7 | -425.69 | 865.83 | 2.11 | 0.21 |

| | | | | | | | | | | |
|---|---|---|---|---|---|---|---|---|---|---|
| -0.03<br>(1.23) | 0.04<br>(0.06) | 0.37<br>(0.19) | 0.07<br>(0.20) | -0.01<br>(0.01) | | 8 | -425.55 | 867.68 | 3.97 | 0.08 |
| 0.42<br>(1.27) | 0.01<br>(0.00) | 0.32<br>(1.89) | -0.01<br>(0.20) | | 0.01<br>(0.30) | 8 | -425.69 | 867.96 | 4.25 | 0.07 |
| -0.02<br>(1.51) | 0.04<br>(0.06) | 0.36<br>(1.89) | 0.06<br>(0.24) | -0.01<br>(0.01) | 0.00<br>(0.30) | 9 | -425.55 | 869.83 | 6.12 | 0.03 |

Gammaglobulin (N = 249 observations, n = 136 individuals)

| I | Age | Pop(CH) | FGM | Age: FGM | Pop(CH): FGM | Mass | df | Log-lik | AICc | delta | AICcw |
|---|---|---|---|---|---|---|---|---|---|---|---|
| **19.88<br>(2.19)** | **0.39<br>(0.12)** | **3.25<br>(1.30)** | | | | **-0.28<br>(0.09)** | **7** | **-647.24** | **1308.94** | **0.00** | **0.56** |
| 21.08<br>(3.47) | 0.38<br>(0.13) | 3.21<br>(1.31) | -0.17<br>(0.39) | | | -0.29<br>(0.09) | 8 | -647.14 | 1310.88 | 1.94 | 0.21 |
| 19.52<br>(4.09) | 0.38<br>(0.13) | 6.59<br>(4.87) | 0.09<br>(0.54) | | -0.54<br>(0.75) | -0.29<br>(0.09) | 9 | -646.88 | 1312.52 | 3.58 | 0.09 |
| 18.14<br>(5.58) | 1.25<br>(1.30) | 3.22<br>(1.31) | 0.27<br>(0.77) | -0.14<br>(0.21) | | -0.28<br>(0.09) | 9 | -646.92 | 1312.59 | 3.65 | 0.09 |
| 17.13<br>(5.82) | 1.14<br>(1.31) | 6.23<br>(4.91) | 0.45<br>(0.82) | -0.12<br>(0.21) | -0.48<br>(0.76) | -0.28<br>(0.10) | 10 | -646.72 | 1314.36 | 5.42 | 0.04 |

Lymphocyte (N = 267 observations, n = 142 individuals)

| I | Age.1 | Age.2 | Pop(CH) | Sex(M) | FGM | Age.1: FGM | Age.2: FGM | Age.2: pop(CH) | Pop(CH): FGM | Sex(M): FGM | Delay | df | Log-lik | AICc | delta | AICcw |
|---|---|---|---|---|---|---|---|---|---|---|---|---|---|---|---|---|
| **2.49<br>(0.81)** | **-0.23<br>(0.08)** | **1.77<br>(0.62)** | **-0.28<br>(0.24)** | **-0.19<br>(0.12)** | **0.19<br>(0.11)** | | **-0.24<br>(0.10)** | **-0.45<br>(0.10)** | | | **-0.00<br>(0.00)** | **12** | **-356.35** | **737.94** | **0.00** | **0.22** |
| 3.12<br>(0.95) | -0.23<br>(0.08) | 1.70<br>(0.63) | -0.27<br>(0.23) | -1.73<br>(1.23) | 0.09<br>(0.14) | | -0.23<br>(0.10) | -0.45<br>(0.10) | | 0.24<br>(0.19) | -0.00<br>(0.00) | 13 | -355.57 | 738.59 | 0.65 | 0.16 |
| 3.79<br>(0.34) | -0.26<br>(0.08) | 0.31<br>(0.06) | -0.30<br>(0.24) | -0.17<br>(0.12) | | | | -0.44<br>(0.10) | | | -0.00<br>(0.00) | 10 | -359.55 | 739.95 | 2.02 | 0.08 |
| 2.28<br>(0.98) | -0.23<br>(0.08) | 1.75<br>(0.63) | 0.21<br>(1.32) | -0.19<br>(0.12) | 0.23<br>(0.14) | | -0.23<br>(0.10) | -0.46<br>(0.10) | -0.08<br>(0.21) | | -0.00<br>(0.00) | 13 | -356.28 | 740.01 | 2.07 | 0.08 |
| 1.87<br>(2.13) | -0.01<br>(0.71) | 1.66<br>(0.71) | -0.28<br>(0.24) | -0.19<br>(0.12) | 0.29<br>(0.33) | -0.04<br>(0.11) | -0.22<br>(0.12) | -0.46<br>(0.10) | | | -0.00<br>(0.00) | 13 | -356.31 | 740.05 | 2.11 | 0.08 |
| 2.84<br>(1.06) | -0.23<br>(0.08) | 1.66<br>(0.63) | 0.51<br>(1.33) | -1.85<br>(1.24) | 0.14<br>(0.16) | | -0.22<br>(0.10) | -0.47<br>(0.10) | -0.12<br>(0.21) | 0.26<br>(0.20) | -0.00<br>(0.00) | 14 | -355.40 | 740.46 | 2.52 | 0.06 |
| 2.62<br>(2.21) | -0.05<br>(0.71) | 1.62<br>(0.71) | -0.27<br>(0.23) | -1.71<br>(1.23) | 0.17<br>(0.35) | -0.03<br>(0.11) | -0.21<br>(0.12) | -0.46<br>(0.10) | | 0.24<br>(0.19) | -0.00<br>(0.00) | 14 | -355.54 | 740.75 | 2.81 | 0.05 |
| 3.11<br>(0.77) | -0.25<br>(0.08) | 0.32<br>(0.06) | -0.27<br>(0.24) | -0.18<br>(0.12) | 0.10<br>(0.10) | | | -0.44<br>(0.10) | | | -0.00<br>(0.00) | 11 | -359.07 | 741.17 | 3.23 | 0.04 |
| 3.81<br>(0.90) | -0.24<br>(0.08) | 0.32<br>(0.06) | -0.26<br>(0.24) | -1.97<br>(1.23) | -0.01<br>(0.13) | | | -0.44<br>(0.10) | | 0.28<br>(0.20) | -0.00<br>(0.00) | 12 | -358.02 | 741.26 | 3.32 | 0.04 |
| 0.53<br>(2.02) | 0.62<br>(0.63) | 0.31<br>(0.06) | -0.28<br>(0.24) | -0.18<br>(0.12) | 0.51<br>(0.31) | -0.14<br>(0.10) | | -0.45<br>(0.10) | | | -0.00<br>(0.00) | 12 | -358.12 | 741.47 | 3.53 | 0.04 |
| 1.39<br>(2.12) | 0.55<br>(0.63) | 0.32<br>(0.06) | -0.27<br>(0.24) | -1.83<br>(1.23) | 0.37<br>(0.33) | -0.13<br>(0.10) | | -0.45<br>(0.10) | | 0.26<br>(0.20) | -0.00<br>(0.00) | 13 | -357.23 | 741.90 | 3.96 | 0.03 |
| 1.67<br>(2.19) | -0.01<br>(0.71) | 1.64<br>(0.71) | 0.21<br>(1.32) | -0.19<br>(0.12) | 0.32<br>(0.34) | -0.03<br>(0.11) | -0.22<br>(0.12) | -0.46<br>(0.10) | -0.08<br>(0.21) | | -0.00<br>(0.00) | 14 | -356.24 | 742.14 | 4.20 | 0.03 |
| 2.36<br>(2.25) | -0.06<br>(0.71) | 1.58<br>(0.71) | 0.50<br>(1.33) | -1.83<br>(1.25) | 0.21<br>(0.35) | -0.03<br>(0.11) | -0.21<br>(0.12) | -0.47<br>(0.10) | -0.12<br>(0.21) | 0.26<br>(0.20) | -0.00<br>(0.00) | 15 | -355.37 | 742.65 | 4.71 | 0.02 |
| 3.36<br>(1.04) | -0.25<br>(0.08) | 0.33<br>(0.06) | 0.87<br>(1.33) | -2.14<br>(1.25) | 0.06<br>(0.15) | | | -0.46<br>(0.10) | -0.18<br>(0.21) | 0.31<br>(0.20) | -0.00<br>(0.00) | 13 | -357.64 | 742.72 | 4.78 | 0.02 |
| 2.74<br>(0.97) | -0.25<br>(0.08) | 0.32<br>(0.06) | 0.54<br>(1.33) | -0.18<br>(0.12) | 0.16<br>(0.14) | | | -0.45<br>(0.11) | -0.13<br>(0.21) | | -0.00<br>(0.00) | 12 | -358.87 | 742.97 | 5.03 | 0.02 |
| 0.27<br>(2.07) | 0.59<br>(0.63) | 0.32<br>(0.06) | 0.43<br>(1.32) | -0.18<br>(0.12) | 0.55<br>(0.32) | -0.13<br>(0.10) | | -0.46<br>(0.10) | -0.11<br>(0.21) | | -0.00<br>(0.00) | 13 | -357.97 | 743.38 | 5.44 | 0.01 |
| 1.11<br>(2.15) | 0.50<br>(0.63) | 0.32<br>(0.06) | 0.74<br>(1.33) | -1.98<br>(1.25) | 0.41<br>(0.33) | -0.12<br>(0.10) | | -0.46<br>(0.10) | -0.16<br>(0.21) | 0.29<br>(0.20) | -0.00<br>(0.00) | 14 | -356.93 | 743.52 | 5.58 | 0.01 |

Gastro-intestinal strongyles (N = 274 observations, n = 140 individuals)

| I | Age.1 | Age.2 | Sex(M) | FGM | Age.1: FGM | Sex(M): FGM | Mass | Age-last | df | Log-lik | AICc | delta | AICcw |
|---|---|---|---|---|---|---|---|---|---|---|---|---|---|
| **3.29 (0.82)** | **0.19 (0.06)** | | **0.56 (0.21)** | | | | **-0.08 (0.04)** | **-0.13 (0.06)** | **8** | **-486.53** | **989.61** | **0.00** | **0.21** |
| 3.25 (0.82) | 0.18 (0.06) | 1.06 (1.33) | 0.56 (0.21) | | | | -0.07 (0.04) | -0.13 (0.06) | 9 | -486.22 | 991.12 | 1.51 | 0.10 |
| 3.25 (0.82) | 0.18 (0.06) | 1.06 (1.33) | 0.56 (0.21) | | | | -0.07 (0.04) | -0.13 (0.06) | 9 | -486.22 | 991.12 | 1.51 | 0.10 |
| 2.56 (1.38) | 0.19 (0.06) | | 0.56 (0.21) | 0.11 (0.17) | | | -0.08 (0.04) | -0.13 (0.06) | 9 | -486.32 | 991.32 | 1.71 | 0.09 |
| 3.39 (1.57) | 0.19 (0.06) | | -1.64 (2.07) | -0.03 (0.22) | | 0.35 (0.33) | -0.07 (0.04) | -0.13 (0.06) | 10 | -485.75 | 992.35 | 2.74 | 0.05 |
| 2.54 (1.37) | 0.18 (0.06) | 1.05 (1.33) | 0.56 (0.21) | 0.11 (0.17) | | | -0.07 (0.04) | -0.13 (0.06) | 10 | -486.01 | 992.86 | 3.25 | 0.04 |
| 2.54 (1.37) | 0.18 (0.06) | 1.05 (1.33) | 0.56 (0.21) | 0.11 (0.17) | | | -0.07 (0.04) | -0.13 (0.06) | 10 | -486.01 | 992.86 | 3.25 | 0.04 |
| 2.54 (1.37) | 0.18 (0.06) | 1.05 (1.33) | 0.56 (0.21) | 0.11 (0.17) | | | -0.07 (0.04) | -0.13 (0.06) | 10 | -486.01 | 992.86 | 3.25 | 0.04 |
| 2.54 (1.37) | 0.18 (0.06) | 1.05 (1.33) | 0.56 (0.21) | 0.11 (0.17) | | | -0.07 (0.04) | -0.13 (0.06) | 10 | -486.01 | 992.86 | 3.25 | 0.04 |
| 3.89 (2.35) | -0.19 (0.55) | | 0.56 (0.21) | -0.09 (0.34) | 0.06 (0.09) | | -0.08 (0.04) | -0.13 (0.06) | 10 | -486.08 | 992.99 | 3.38 | 0.04 |
| 3.37 (1.57) | 0.19 (0.06) | 1.07 (1.34) | -1.65 (2.06) | -0.03 (0.22) | | 0.35 (0.32) | -0.07 (0.04) | -0.13 (0.06) | 11 | -485.44 | 993.88 | 4.28 | 0.02 |
| 3.37 (1.57) | 0.19 (0.06) | 1.07 (1.34) | -1.65 (2.06) | -0.03 (0.22) | | 0.35 (0.32) | -0.07 (0.04) | -0.13 (0.06) | 11 | -485.44 | 993.88 | 4.28 | 0.02 |
| 3.37 (1.57) | 0.19 (0.06) | 1.07 (1.34) | -1.65 (2.06) | -0.03 (0.22) | | 0.35 (0.32) | -0.07 (0.04) | -0.13 (0.06) | 11 | -485.44 | 993.88 | 4.28 | 0.02 |
| 3.37 (1.57) | 0.19 (0.06) | 1.07 (1.34) | -1.65 (2.06) | -0.03 (0.22) | | 0.35 (0.32) | -0.07 (0.04) | -0.13 (0.06) | 11 | -485.44 | 993.88 | 4.28 | 0.02 |
| 4.87 (2.51) | -0.22 (0.55) | | -1.73 (2.07) | -0.26 (0.37) | 0.07 (0.09) | 0.36 (0.33) | -0.08 (0.04) | -0.13 (0.06) | 11 | -485.47 | 993.94 | 4.33 | 0.02 |
| 3.88 (2.35) | -0.20 (0.55) | 1.06 (1.33) | 0.56 (0.21) | -0.10 (0.34) | 0.06 (0.09) | | -0.08 (0.04) | -0.13 (0.06) | 11 | -485.76 | 994.54 | 4.93 | 0.02 |
| 3.88 (2.35) | -0.20 (0.55) | 1.06 (1.33) | 0.56 (0.21) | -0.10 (0.34) | 0.06 (0.09) | | -0.08 (0.04) | -0.13 (0.06) | 11 | -485.76 | 994.54 | 4.93 | 0.02 |
| 3.88 (2.35) | -0.20 (0.55) | 1.06 (1.33) | 0.56 (0.21) | -0.10 (0.34) | 0.06 (0.09) | | -0.08 (0.04) | -0.13 (0.06) | 11 | -485.76 | 994.54 | 4.93 | 0.02 |
| 3.88 (2.35) | -0.20 (0.55) | 1.06 (1.33) | 0.56 (0.21) | -0.10 (0.34) | 0.06 (0.09) | | -0.08 (0.04) | -0.13 (0.06) | 11 | -485.76 | 994.54 | 4.93 | 0.02 |
| 4.87 (2.51) | -0.23 (0.55) | 1.08 (1.33) | -1.74 (2.06) | -0.27 (0.37) | 0.07 (0.09) | 0.36 (0.33) | -0.07 (0.04) | -0.13 (0.06) | 12 | -485.14 | 995.48 | 5.87 | 0.01 |
| 4.87 (2.51) | -0.23 (0.55) | 1.08 (1.33) | -1.74 (2.06) | -0.27 (0.37) | 0.07 (0.09) | 0.36 (0.33) | -0.07 (0.04) | -0.13 (0.06) | 12 | -485.14 | 995.48 | 5.87 | 0.01 |
| 4.87 (2.51) | -0.23 (0.55) | 1.08 (1.33) | -1.74 (2.06) | -0.27 (0.37) | 0.07 (0.09) | 0.36 (0.33) | -0.07 (0.04) | -0.13 (0.06) | 12 | -485.14 | 995.48 | 5.87 | 0.01 |
| 4.87 (2.51) | -0.23 (0.55) | 1.08 (1.33) | -1.74 (2.06) | -0.27 (0.37) | 0.07 (0.09) | 0.36 (0.33) | -0.07 (0.04) | -0.13 (0.06) | 12 | -485.14 | 995.48 | 5.87 | 0.01 |

*Trichuris* sp. (N = 252 observations, n = 129 individuals)

| I | Age.1 | Age.2 | Pop(CH) | Sex(M) | FGM | Age.1: pop(CH) | Age.1: sex(M) | Age.1: FGM | Pop(CH): FGM | Sex(M): FGM | Mass | df | Log-lik | AICc | delta | AICcw |
|---|---|---|---|---|---|---|---|---|---|---|---|---|---|---|---|---|
| **2.00 (0.82)** | **-0.06 (0.08)** | **4.40 (1.34)** | **0.45 (0.40)** | **0.19 (0.38)** | | **0.11 (0.10)** | **0.10 (0.10)** | | | | **-0.08 (0.04)** | **11** | **-420.17** | **863.44** | **0.00** | **0.33** |
| 2.49 (1.28) | -0.07 (0.08) | 4.40 (1.34) | 0.43 (0.40) | 0.20 (0.38) | -0.07 (0.15) | 0.11 (0.10) | 0.10 (0.10) | | | | -0.08 (0.04) | 12 | -420.05 | 865.42 | 1.97 | 0.13 |
| 2.49 (1.28) | -0.07 (0.08) | 4.40 (1.34) | 0.43 (0.40) | 0.20 (0.38) | -0.07 (0.15) | 0.11 (0.10) | 0.10 (0.10) | | | | -0.08 (0.04) | 12 | -420.05 | 865.42 | 1.97 | 0.13 |
| 1.93 (1.44) | -0.06 (0.08) | 4.38 (1.34) | 0.40 (0.41) | 1.88 (2.01) | 0.03 (0.19) | 0.11 (0.10) | 0.09 (0.10) | | | -0.26 (0.31) | -0.08 (0.04) | 13 | -419.69 | 866.92 | 3.47 | 0.06 |

| | | | | | | | | | | | | | | | | | | | | | |
|---|---|---|---|---|---|---|---|---|---|---|---|---|---|---|---|---|---|---|---|---|---|
| 1.93 (1.44) | -0.06 (0.08) | 4.38 (1.34) | 0.40 (0.41) | 1.88 (2.01) | 0.03 (0.19) | 0.11 (0.10) | 0.09 (0.10) | | | -0.26 (0.31) | -0.08 (0.04) | | | | | | 13 | -419.69 | 866.92 | 3.47 | 0.06 |
| 2.15 (1.55) | -0.07 (0.08) | 4.40 (1.34) | 1.21 (2.00) | 0.19 (0.38) | -0.02 (0.21) | 0.10 (0.10) | 0.10 (0.10) | | -0.12 (0.31) | | -0.08 (0.04) | | | | | | 13 | -419.98 | 867.49 | 4.04 | 0.04 |
| 2.15 (1.55) | -0.07 (0.08) | 4.40 (1.34) | 1.21 (2.00) | 0.19 (0.38) | -0.02 (0.21) | 0.10 (0.10) | 0.10 (0.10) | | -0.12 (0.31) | | -0.08 (0.04) | | | | | | 13 | -419.98 | 867.49 | 4.04 | 0.04 |
| 3.06 (2.33) | -0.23 (0.57) | 4.40 (1.34) | 0.42 (0.41) | 0.19 (0.38) | -0.16 (0.33) | 0.12 (0.10) | 0.10 (0.10) | 0.03 (0.09) | | | -0.08 (0.04) | | | | | | 13 | -420.01 | 867.56 | 4.11 | 0.04 |
| 3.06 (2.33) | -0.23 (0.57) | 4.40 (1.34) | 0.42 (0.41) | 0.19 (0.38) | -0.16 (0.33) | 0.12 (0.10) | 0.10 (0.10) | 0.03 (0.09) | | | -0.08 (0.04) | | | | | | 13 | -420.01 | 867.56 | 4.11 | 0.04 |
| 1.71 (1.64) | -0.06 (0.08) | 4.38 (1.34) | 0.97 (2.02) | 1.80 (2.03) | 0.06 (0.23) | 0.11 (0.10) | 0.09 (0.10) | | -0.09 (0.31) | -0.25 (0.31) | -0.08 (0.04) | | | | | | 14 | -419.65 | 869.08 | 5.64 | 0.02 |
| 1.71 (1.64) | -0.06 (0.08) | 4.38 (1.34) | 0.97 (2.02) | 1.80 (2.03) | 0.06 (0.23) | 0.11 (0.10) | 0.09 (0.10) | | -0.09 (0.31) | -0.25 (0.31) | -0.08 (0.04) | | | | | | 14 | -419.65 | 869.08 | 5.64 | 0.02 |
| 2.28 (2.51) | -0.16 (0.57) | 4.38 (1.34) | 0.39 (0.41) | 1.83 (2.03) | -0.03 (0.37) | 0.12 (0.10) | 0.09 (0.10) | 0.02 (0.09) | | -0.25 (0.31) | -0.08 (0.04) | | | | | | 14 | -419.68 | 869.13 | 5.69 | 0.02 |
| 2.28 (2.51) | -0.16 (0.57) | 4.38 (1.34) | 0.39 (0.41) | 1.83 (2.03) | -0.03 (0.37) | 0.12 (0.10) | 0.09 (0.10) | 0.02 (0.09) | | -0.25 (0.31) | -0.08 (0.04) | | | | | | 14 | -419.68 | 869.13 | 5.69 | 0.02 |
| 2.75 (2.45) | -0.24 (0.57) | 4.40 (1.33) | 1.23 (2.00) | 0.18 (0.38) | -0.11 (0.35) | 0.11 (0.10) | 0.11 (0.10) | 0.03 (0.09) | -0.13 (0.31) | | -0.08 (0.04) | | | | | | 14 | -419.93 | 869.64 | 6.19 | 0.02 |
| 2.75 (2.45) | -0.24 (0.57) | 4.40 (1.33) | 1.23 (2.00) | 0.18 (0.38) | -0.11 (0.35) | 0.11 (0.10) | 0.11 (0.10) | 0.03 (0.09) | -0.13 (0.31) | | -0.08 (0.04) | | | | | | 14 | -419.93 | 869.64 | 6.19 | 0.02 |
| 2.10 (2.59) | -0.17 (0.58) | 4.38 (1.33) | 0.99 (2.02) | 1.74 (2.06) | 0.00 (0.38) | 0.11 (0.10) | 0.09 (0.10) | 0.02 (0.09) | -0.09 (0.31) | -0.24 (0.31) | -0.08 (0.04) | | | | | | 15 | -419.64 | 871.31 | 7.86 | 0.01 |
| 2.10 (2.59) | -0.17 (0.58) | 4.38 (1.33) | 0.99 (2.02) | 1.74 (2.06) | 0.00 (0.38) | 0.11 (0.10) | 0.09 (0.10) | 0.02 (0.09) | -0.09 (0.31) | -0.24 (0.31) | -0.08 (0.04) | | | | | | 15 | -419.64 | 871.31 | 7.86 | 0.01 |

Protostrongylids (N = 256 observations, n = 133 individuals)

| I | Age | Age² | FGM | Pop(CH) | Sex(M) | Cohort(P) | Age: cohort(P) | Age: FGM | Age²: sex(M) | Age²: cohort(P) | Age²: FGM | Pop(CH): FGM | Sex(M): FGM | Cohort(P): FGM | Mass | Age-last | df | Log-lik | AICc | delta | AICcw |
|---|---|---|---|---|---|---|---|---|---|---|---|---|---|---|---|---|---|---|---|---|---|
| 0.27 (1.06) | -0.04 (0.14) | 0.01 (0.01) | 0.10 (0.15) | 2.10 (1.09) | -1.68 (1.16) | -2.83 (1.21) | 0.04 (0.23) | | 0.01 (0.01) | 0.00 (0.02) | | -0.34 (0.17) | 0.29 (0.18) | 0.45 (0.18) | -0.04 (0.02) | 0.02 (0.03) | 18 | -280.65 | 600.16 | 0.00 | 0.12 |
| -0.12 (1.05) | -0.05 (0.14) | 0.01 (0.01) | 0.17 (0.14) | 1.87 (1.10) | 0.14 (0.15) | -3.31 (1.20) | 0.07 (0.23) | | 0.01 (0.01) | 0.00 (0.02) | | -0.31 (0.17) | | 0.51 (0.18) | -0.05 (0.02) | 0.02 (0.03) | 17 | -281.84 | 600.24 | 0.07 | 0.11 |
| 1.85 (1.52) | -0.51 (0.35) | 0.01 (0.01) | -0.14 (0.22) | 2.26 (1.07) | -1.95 (1.15) | -2.69 (1.20) | 0.01 (0.23) | 0.08 (0.05) | 0.01 (0.01) | 0.01 (0.02) | | -0.37 (0.17) | 0.33 (0.18) | 0.43 (0.17) | -0.05 (0.02) | 0.02 (0.03) | 19 | -279.64 | 600.47 | 0.30 | 0.10 |
| **0.71 (0.95)** | **-0.03 (0.14)** | **0.01 (0.01)** | **0.04 (0.12)** | **-0.06 (0.13)** | **0.15 (0.15)** | **-3.24 (1.21)** | **0.08 (0.23)** | | **0.01 (0.01)** | **0.00 (0.02)** | | | | **0.50 (0.18)** | **-0.05 (0.02)** | **0.02 (0.03)** | **16** | **-283.36** | **600.97** | **0.81** | **0.08** |
| 0.92 (1.18) | -0.05 (0.14) | -0.04 (0.04) | 0.00 (0.17) | 2.23 (1.08) | -1.91 (1.16) | -2.69 (1.21) | 0.00 (0.23) | | 0.01 (0.01) | 0.01 (0.03) | 0.01 (0.01) | -0.36 (0.17) | 0.32 (0.18) | 0.43 (0.18) | -0.05 (0.02) | 0.02 (0.03) | 19 | -279.93 | 601.05 | 0.88 | 0.07 |
| 1.07 (1.48) | -0.41 (0.34) | 0.01 (0.01) | -0.01 (0.21) | 1.96 (1.09) | 0.14 (0.14) | -3.26 (1.19) | 0.05 (0.23) | 0.06 (0.05) | 0.01 (0.01) | 0.01 (0.02) | | -0.32 (0.17) | | 0.51 (0.17) | -0.05 (0.02) | 0.02 (0.03) | 18 | -281.21 | 601.29 | 1.12 | 0.07 |
| 1.11 (0.99) | -0.02 (0.14) | 0.01 (0.01) | -0.04 (0.13) | -0.03 (0.13) | -1.35 (1.16) | -2.84 (1.23) | 0.05 (0.22) | | 0.01 (0.01) | 0.00 (0.02) | | | 0.24 (0.18) | 0.44 (0.18) | -0.04 (0.02) | 0.02 (0.03) | 17 | -282.54 | 601.63 | 1.46 | 0.06 |
| 0.34 (1.16) | -0.05 (0.14) | -0.03 (0.04) | 0.11 (0.16) | 1.94 (1.10) | 0.14 (0.15) | -3.26 (1.20) | 0.05 (0.23) | | 0.01 (0.01) | 0.01 (0.03) | 0.01 (0.01) | -0.32 (0.17) | | 0.51 (0.17) | -0.05 (0.02) | 0.02 (0.03) | 18 | -281.43 | 601.72 | 1.55 | 0.05 |
| 4.07 (2.87) | -1.81 (1.47) | 0.17 (0.18) | -0.50 (0.45) | 2.26 (1.06) | -1.94 (1.14) | -2.75 (1.20) | 0.05 (0.23) | 0.29 (0.24) | 0.01 (0.01) | 0.00 (0.03) | -0.03 (0.03) | -0.36 (0.17) | 0.33 (0.18) | 0.43 (0.17) | -0.05 (0.02) | 0.02 (0.03) | 20 | -279.25 | 602.04 | 1.87 | 0.05 |
| 1.75 (1.44) | -0.33 (0.34) | 0.01 (0.01) | -0.13 (0.21) | -0.05 (0.13) | 0.16 (0.15) | -3.19 (1.20) | 0.06 (0.23) | 0.05 (0.05) | 0.01 (0.01) | 0.01 (0.02) | | | | 0.50 (0.18) | -0.05 (0.02) | 0.02 (0.03) | 17 | -282.90 | 602.36 | 2.19 | 0.04 |
| 2.46 (1.51) | -0.40 (0.35) | 0.01 (0.01) | -0.25 (0.22) | -0.03 (0.13) | -1.55 (1.16) | -2.74 (1.22) | 0.03 (0.23) | 0.06 (0.05) | 0.01 (0.01) | 0.01 (0.02) | | | 0.27 (0.18) | 0.43 (0.18) | -0.05 (0.02) | 0.02 (0.03) | 18 | -281.86 | 602.57 | 2.41 | 0.03 |
| 1.11 (1.09) | -0.03 (0.14) | -0.02 (0.04) | -0.02 (0.14) | -0.06 (0.13) | 0.16 (0.15) | -3.19 (1.21) | 0.06 (0.23) | | 0.00 (0.01) | 0.01 (0.03) | 0.00 (0.01) | | | 0.50 (0.18) | -0.05 (0.02) | 0.02 (0.03) | 17 | -283.07 | 602.70 | 2.53 | 0.03 |
| 3.30 (2.87) | -1.72 (1.47) | 0.17 (0.18) | -0.37 (0.45) | 1.96 (1.08) | 0.14 (0.14) | -3.32 (1.19) | 0.09 (0.23) | 0.27 (0.24) | 0.01 (0.01) | 0.00 (0.03) | -0.03 (0.03) | -0.32 (0.17) | | 0.51 (0.17) | -0.05 (0.02) | 0.02 (0.03) | 19 | -280.82 | 602.84 | 2.68 | 0.03 |
| 1.68 (1.15) | -0.02 (0.14) | -0.03 (0.04) | -0.12 (0.16) | -0.03 (0.13) | -1.51 (1.16) | -2.75 (1.22) | 0.03 (0.23) | | 0.01 (0.01) | 0.01 (0.03) | 0.01 (0.01) | | 0.26 (0.18) | 0.43 (0.18) | -0.05 (0.02) | 0.02 (0.03) | 18 | -282.08 | 603.02 | 2.85 | 0.03 |
| 3.91 (2.85) | -1.60 (1.48) | 0.16 (0.18) | -0.48 (0.45) | -0.05 (0.13) | 0.15 (0.15) | -3.26 (1.20) | 0.10 (0.23) | 0.26 (0.24) | 0.01 (0.01) | -0.00 (0.03) | -0.03 (0.03) | | | 0.50 (0.17) | -0.05 (0.02) | 0.02 (0.03) | 18 | -282.54 | 603.94 | 3.77 | 0.02 |

| | | | | | | | | | | | | | | | | | | | | | |
|---|---|---|---|---|---|---|---|---|---|---|---|---|---|---|---|---|---|---|---|---|---|
| 1.06 (1.50) | -0.52 (0.35) | 0.01 (0.01) | -0.02 (0.22) | 2.25 (1.09) | -2.61 (1.14) | 0.05 (0.46) | -0.01 (0.23) | 0.08 (0.05) | 0.01 (0.01) | 0.01 (0.03) | | -0.37 (0.17) | 0.44 (0.18) | | -0.05 (0.02) | 0.02 (0.03) | 18 | -282.58 | 604.03 | 3.87 | 0.02 |
| -0.70 (1.00) | -0.02 (0.14) | 0.01 (0.01) | 0.26 (0.14) | 2.06 (1.11) | -2.33 (1.15) | 0.02 (0.46) | 0.02 (0.23) | | 0.01 (0.01) | 0.00 (0.02) | | -0.34 (0.18) | 0.39 (0.18) | | -0.05 (0.02) | 0.02 (0.03) | 17 | -283.74 | 604.04 | 3.87 | 0.02 |
| 4.59 (2.87) | -1.66 (1.48) | 0.16 (0.18) | -0.59 (0.45) | -0.02 (0.12) | -1.54 (1.15) | -2.79 (1.21) | 0.07 (0.23) | 0.27 (0.24) | 0.01 (0.01) | 0.00 (0.03) | -0.03 (0.03) | | 0.27 (0.18) | 0.43 (0.18) | -0.05 (0.02) | 0.02 (0.03) | 19 | -281.49 | 604.18 | 4.01 | 0.02 |
| 0.06 (1.15) | -0.02 (0.14) | -0.05 (0.04) | 0.14 (0.16) | 2.22 (1.10) | -2.56 (1.14) | 0.07 (0.46) | -0.01 (0.23) | | 0.01 (0.01) | 0.01 (0.03) | 0.01 (0.01) | -0.36 (0.17) | 0.43 (0.18) | | -0.05 (0.02) | 0.02 (0.03) | 18 | -282.89 | 604.64 | 4.48 | 0.01 |
| 0.13 (0.92) | -0.00 (0.14) | 0.00 (0.01) | 0.12 (0.12) | -0.05 (0.13) | -1.96 (1.16) | -0.03 (0.46) | 0.05 (0.23) | | 0.01 (0.01) | 0.00 (0.02) | | | 0.34 (0.18) | | -0.05 (0.02) | 0.02 (0.03) | 16 | -285.53 | 605.31 | 5.15 | 0.01 |
| 3.19 (2.88) | -1.76 (1.49) | 0.16 (0.18) | -0.36 (0.45) | 2.24 (1.08) | -2.60 (1.13) | -0.00 (0.46) | 0.03 (0.23) | 0.28 (0.24) | 0.01 (0.01) | 0.00 (0.03) | -0.03 (0.03) | -0.36 (0.17) | 0.44 (0.18) | | -0.05 (0.02) | 0.02 (0.03) | 19 | -282.23 | 605.66 | 5.50 | 0.01 |
| 1.64 (1.49) | -0.41 (0.35) | 0.00 (0.01) | -0.11 (0.22) | -0.05 (0.13) | -2.16 (1.15) | -0.01 (0.46) | 0.03 (0.23) | 0.07 (0.05) | 0.01 (0.01) | 0.00 (0.03) | | | 0.37 (0.18) | | -0.05 (0.02) | 0.02 (0.03) | 17 | -284.73 | 606.01 | 5.85 | 0.01 |
| -1.48 (0.96) | -0.01 (0.14) | 0.01 (0.01) | 0.40 (0.12) | 1.69 (1.14) | 0.17 (0.15) | -0.02 (0.46) | 0.06 (0.23) | | 0.01 (0.01) | 0.00 (0.02) | | -0.28 (0.18) | | | -0.05 (0.02) | 0.02 (0.03) | 16 | -285.98 | 606.21 | 6.05 | 0.01 |
| -0.67 (0.83) | 0.00 (0.14) | 0.00 (0.01) | 0.26 (0.09) | -0.09 (0.13) | 0.19 (0.15) | -0.05 (0.46) | 0.07 (0.23) | | 0.01 (0.01) | 0.00 (0.02) | | | | | -0.05 (0.02) | 0.02 (0.03) | 15 | -287.19 | 606.36 | 6.19 | 0.01 |
| 0.79 (1.11) | -0.00 (0.14) | -0.04 (0.04) | 0.02 (0.15) | -0.05 (0.13) | -2.13 (1.15) | 0.01 (0.46) | 0.02 (0.23) | | 0.01 (0.01) | 0.01 (0.03) | 0.01 (0.01) | | 0.36 (0.18) | | -0.05 (0.02) | 0.02 (0.03) | 17 | -284.96 | 606.48 | 6.31 | 0.00 |
| -0.28 (1.44) | -0.38 (0.35) | 0.01 (0.01) | 0.21 (0.21) | 1.80 (1.13) | 0.18 (0.15) | -0.01 (0.47) | 0.04 (0.23) | 0.06 (0.05) | 0.01 (0.01) | 0.00 (0.03) | | -0.30 (0.18) | | | -0.05 (0.02) | 0.02 (0.03) | 17 | -285.35 | 607.26 | 7.09 | 0.00 |
| -1.01 (1.09) | -0.02 (0.14) | -0.03 (0.04) | 0.33 (0.15) | 1.77 (1.13) | 0.18 (0.15) | 0.00 (0.47) | 0.04 (0.23) | | 0.01 (0.01) | 0.00 (0.03) | 0.01 (0.01) | -0.30 (0.18) | | | -0.05 (0.02) | 0.02 (0.03) | 17 | -285.55 | 607.65 | 7.49 | 0.00 |
| 3.69 (2.88) | -1.62 (1.49) | 0.15 (0.18) | -0.45 (0.45) | -0.04 (0.13) | -2.16 (1.14) | -0.06 (0.47) | 0.06 (0.23) | 0.26 (0.24) | 0.01 (0.01) | -0.00 (0.03) | -0.02 (0.03) | | 0.37 (0.18) | | -0.05 (0.02) | 0.02 (0.03) | 18 | -284.40 | 607.67 | 7.51 | 0.00 |
| 0.40 (1.39) | -0.31 (0.35) | 0.00 (0.01) | 0.10 (0.19) | -0.09 (0.13) | 0.19 (0.15) | -0.03 (0.46) | 0.05 (0.23) | 0.05 (0.05) | 0.01 (0.01) | 0.00 (0.02) | | | | | -0.05 (0.02) | 0.02 (0.03) | 16 | -286.72 | 607.70 | 7.54 | 0.00 |
| -0.24 (1.00) | 0.00 (0.14) | -0.03 (0.04) | 0.20 (0.12) | -0.09 (0.13) | 0.19 (0.15) | -0.02 (0.46) | 0.05 (0.23) | | 0.01 (0.01) | 0.00 (0.03) | 0.00 (0.01) | | | | -0.05 (0.02) | 0.02 (0.03) | 16 | -286.88 | 608.03 | 7.86 | 0.00 |
| 1.78 (2.87) | -1.58 (1.49) | 0.16 (0.18) | -0.12 (0.45) | 1.80 (1.12) | 0.17 (0.15) | -0.07 (0.47) | 0.08 (0.23) | 0.25 (0.24) | 0.01 (0.01) | -0.00 (0.03) | -0.02 (0.03) | -0.30 (0.18) | | | -0.06 (0.02) | 0.02 (0.03) | 18 | -285.03 | 608.93 | 8.76 | 0.00 |
| 2.38 (2.84) | -1.46 (1.49) | 0.15 (0.18) | -0.22 (0.44) | -0.08 (0.13) | 0.19 (0.15) | -0.09 (0.47) | 0.09 (0.23) | 0.24 (0.24) | 0.01 (0.01) | -0.00 (0.03) | -0.02 (0.03) | | | | -0.06 (0.02) | 0.02 (0.03) | 17 | -286.42 | 609.40 | 9.23 | 0.00 |
| 1.15 (0.54) | -0.02 (0.14) | 0.01 (0.01) | | -0.15 (0.14) | 0.19 (0.15) | -0.12 (0.47) | 0.11 (0.23) | | 0.01 (0.01) | -0.00 (0.02) | | | | | -0.05 (0.02) | 0.01 (0.03) | 14 | -291.04 | 611.80 | 11.64 | 0.00 |

Coccidia (N = 272 observations, n = 140 individuals)

| I | FGM | df | Log-lik | AICc | delta | AICcw |
|---|---|---|---|---|---|---|
| **1.25 (0.12)** | | **4** | **-568.93** | **1146.01** | **0.00** | **0.52** |
| -0.46 (1.24) | 0.27 (0.20) | 5 | -567.97 | 1146.16 | 0.15 | 0.48 |
